# Control of working memory maintenance by theta-gamma phase amplitude coupling of human hippocampal neurons

**DOI:** 10.1101/2023.04.05.535772

**Authors:** Jonathan Daume, Jan Kaminski, Andrea G. P. Schjetnan, Yousef Salimpour, Umais Khan, Chrystal Reed, William Anderson, Taufik A. Valiante, Adam N. Mamelak, Ueli Rutishauser

## Abstract

Retaining information in working memory (WM) is a demanding process that relies on cognitive control to protect memoranda-specific persistent activity from interference. How cognitive control regulates WM storage, however, remains unknown. We hypothesized that interactions of frontal control and hippocampal persistent activity are coordinated by theta-gamma phase amplitude coupling (TG-PAC). We recorded single neurons in the human medial temporal and frontal lobe while patients maintained multiple items in WM. In the hippocampus, TG-PAC was indicative of WM load and quality. We identified cells that selectively spiked during nonlinear interactions of theta phase and gamma amplitude. These PAC neurons were more strongly coordinated with frontal theta activity when cognitive control demand was high, and they introduced information-enhancing and behaviorally relevant noise correlations with persistently active neurons in the hippocampus. We show that TG-PAC integrates cognitive control and WM storage to improve the fidelity of WM representations and facilitate behavior.

## Introduction

Working memory (WM), the ability to actively maintain and manipulate a limited amount of information in mind for a brief period of time (Baddeley 1992), is a crucial component of cognition. Deficits in WM due to disease, age, developmental disruptions, fatigue, or cognitive overload can have serious detrimental effects on neurotypical cognitive function. It is therefore of critical importance to develop a better understanding of the underlying mechanisms which remain far from understood.

WM maintenance is an active process that retains information that is no longer available in the external world. One cellular mechanism that is thought to support this process is persistent neural activity (Fuster and Alexander 1971; Constantinidis and Klingberg 2016; Leavitt et al. 2017a; Zylberberg and Strowbridge 2017; Kamiński and Rutishauser 2020; Wang 2021). In humans, memoranda-specific persistent activity has been observed in cells of the human medial temporal lobe (MTL) but not frontal lobe (Kamiński et al. 2017; Kornblith et al. 2017; Boran et al. 2019). Accurately maintaining persistent activity is demanding and must resist interference introduced by distractors or high memory load. These are the conditions under which the MTL, in addition to its classic role in long-term memory, also becomes essential for short-term memories such as WM (Jeneson and Squire 2012). It is thought that cognitive control (variously referred to as top-down, executive, or attentional control by others), is needed to support the active maintenance of WM content (Cowan 2010; Baddeley 2012). Classic abstract cognitive models of WM assign the role of control to the ‘central executive’, suggested to be a function of the frontal lobes (Baddeley 2003; Badre 2008; Helfrich and Knight 2016; Badre and Nee 2018), and its interactions with temporal and parietal storage systems to support WM maintenance (Curtis and D’Esposito 2003; D’Esposito and Postle 2015; Lara and Wallis 2015; Scimeca et al. 2018). However, at the implementation level, little is known about how cognitive control and storage mechanisms interact.

A ubiquitous macroscopic phenomenon that has been associated with both WM maintenance and frontal control over WM content-specific processing is theta-gamma phase amplitude coupling (PAC) (Lisman and Jensen 2013). PAC has been documented in a large number of tasks and brain areas at the level of the local field potential (LFP). Despite its ubiquity, however, the functional role of these interactions remains largely unknown. A major hypothesis that has been advanced in recent years is that cross-frequency interactions such as PAC enable the integration of local sensory information processing with brain-wide cognitive control (Canolty and Knight 2010; Palva and Palva 2018). Sensory processing leads to local increases in power in the gamma range (30– 140 Hz) (Colgin et al. 2009; Yamamoto et al. 2014; Fries 2015; Colgin 2016; Fernández-Ruiz et al. 2021), whereas cognitive control regulates distant brain processes through long-range inter-areal interactions in the theta range (3-7 Hz) (Miller 2000; Fell and Axmacher 2011; Cavanagh and Frank 2014; Harris and Gordon 2015). Under this framework, theta-gamma PAC serves as a tool to integrate these two processes. Supporting this idea, during WM maintenance, PAC has been observed in areas of the temporal lobe, in particular the hippocampus (Canolty et al. 2006; Axmacher et al. 2010; Yamamoto et al. 2014; Leszczyński et al. 2015; Reinhart and Nguyen 2019; Abubaker et al. 2021), where it co-occurs with long-range theta phase synchronization to frontal regions and thus reflects the theta-based frontal coordination of high-frequency WM content processing in temporal areas (Daume et al. 2017b, a). Here, we test the hypothesis that neurons in the human MTL whose activity is modulated by both theta phase and gamma amplitude in ways predicted by PAC exist and that these neurons enable PAC-mediated inter-areal interactions that enhance the ability to maintain WM content by persistently active neurons.

To study this question, we recorded single cell activity as well as LFPs from medial frontal and MTL areas while human neurosurgical patients performed a Sternberg WM task. We developed a new method to identify “PAC neurons”, which were common in areas of the MTL. PAC neurons were not tuned to WM content. In contrast, WM-selective category neurons remained persistently active when their preferred picture category was maintained in WM. Hippocampal PAC neurons coordinated their activity with frontal theta activity in trials with higher WM load and faster reaction times, indicating that they are related to cognitive control processes. PAC neurons shaped the population-level geometry of WM representations and enhanced WM fidelity by introducing noise correlations with content-tuned category neurons in the hippocampus. Taken together, we show that PAC neurons have a functional role in inter-areal cognitive control of WM content-specific persistent activity within the hippocampus.

## Results

### Task, behavioral results, and electrophysiology

36 patients (44 sessions; Table S4) participated in a modified Sternberg WM task with pictures as stimuli. All pictures belonged to one of five different categories (people, animals, cars/tools (depending on variant), fruit, and landscapes). In each trial, patients were asked to maintain either one (load 1) or three (load 3) consecutively presented pictures in their WM for 2.5-2.8 s (Fig. 1a). Following the maintenance period, patients were asked whether the probe stimulus shown was identical to one of the item(s) they were holding in WM. Patients performed well, with a mean accuracy of 93.66 ± 7.04 % (78.34 ± 20.09 % of all errors were false negative). Across all sessions, subjects responded slower (1.46 s vs 1.33 s; t(43) = 6.42; p < 0.001) and less accurate (91.60 % vs 95.71 %; t(43) = -4.45; p < 0.001) in load 3 as compared to load 1 trials (Fig. 1d). While patients performed the task, we recorded in total from 1452 single neurons across five brain regions. 360 neurons were located in the hippocampus, 496 in the amygdala, 204 in the pre-supplementary motor area (pre-SMA), 188 in the dorsal anterior cingulate cortex (dACC), and 206 in the ventromedial prefrontal cortex (vmPFC; Fig. 1b, c). At the same time, we also recorded the broadband LFP from a total of 1911 microwires (a subset of which provided the single neuron data). Of the LFP channels, 586 channels were in the hippocampus, 416 in the amygdala, 283 in the pre-SMA, 319 in the dACC, and 307 in the vmPFC.

**Figure 1.**
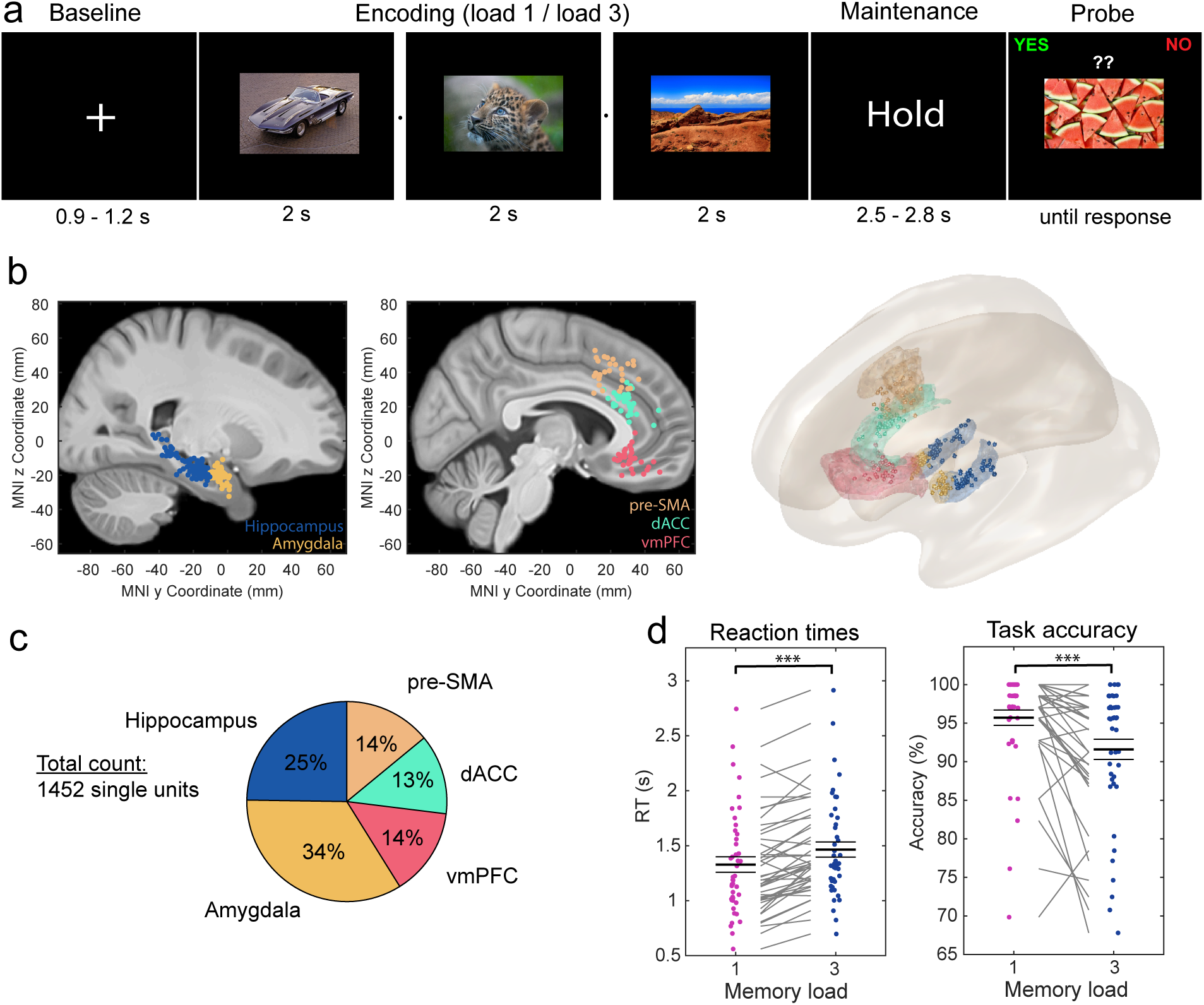
Task, recordings sites, and behavior. **(a)** Example trial. Each trial began with a fixation cross followed by either one (load 1) and three (load 3) consecutively presented pictures, each presented for 2 s (separated by a variable blank screen of up to 200 ms as indicated by a small dot). After a variable maintenance period of on average 2.7 s duration, a probe picture was presented. The task was to decide whether the probe picture has been part of the pictures shown during encoding in this trial (correct answer “No” in the example shown). Pictures were drawn from five categories: people, animals, cars/tools, fruit, landscapes. **(b)** Recording locations. Each colored dots represent the location of a micro-wire bundle across all 44 sessions shown on a standardized MNI152 brain template (left) and in a 3D model using the brainnetome atlas (right). **(c)** Proportions of neurons recorded in each brain area. The three frontal areas (pre-SMA, dACC, vmPFC) are jointly referred to as medial frontal cortex (MFC). **(d)** Behavioral results. Patients made fewer errors and responded faster in load 1 as compared to load 3 trials. Each line connects the two dots belonging to the same session. RT was measured relative to probe stimulus onset. *** p < 0.001, permutation-based t-test.

### Theta-gamma PAC differed as a function of WM load and correlated with reaction times in the hippocampus

We first determined whether we observed PAC at the level of the LFP during WM maintenance and if so whether it differed as a function of WM load. For that purpose, we estimated PAC for the LFP recorded on all micro wires using a modulation index that varied as a function of low frequency phase and high frequency power. We used all correct trials from both load conditions to do so. The raw modulations indices from each channel and frequency combination were normalized using a surrogate distribution to derive a z-scored version (using 200 random shuffles, see Methods). Averaging the normalized modulation indices across all recording channels (Fig. 2a) revealed that the strongest PAC was between the phase of the LFP in the theta range (3-7 Hz) and the amplitude in two different gamma frequency bands, a lower (30-55 Hz) and a higher gamma range (70-140 Hz), consistent with what has been found in earlier intracranial recordings from humans and rodents (Canolty et al. 2006; Colgin et al. 2009; Yamamoto et al. 2014). For each channel, we then separately averaged the normalized modulation indices in the theta to low gamma and the theta to high gamma combinations and assessed which of the channels exhibited significant PAC across both load conditions in the given frequency combination (averaged z-score > 1.64; p < 0.05; right-sided). The selected channels were then used to compare the within-condition normalized PAC estimates between the two load conditions in each brain area. For theta to high gamma PAC, 137 (23.38 %) out of a total of 586 hippocampal channels showed significant PAC across all correct trials from both load conditions. Comparing the PAC estimates between load conditions in the selected channels revealed significantly weaker theta-high gamma PAC in load 3 compared to load 1 trials (t(136) = -4.26, p < 0.001; FDR corrected for the five brain regions of interest (Benjamini and Hochberg 1995); Fig. 2c shows the comodulogram for significant PAC channels from the hippocampus per load condition). In the amygdala, a substantial proportion of channels exhibited significant PAC across both conditions (130 (31.25%) out of 416 possible channels). However, in contrast to the hippocampus, theta-high gamma PAC did not differ significantly between the two load conditions in the amygdala (t(129) = 1.43, p = 0.38). Furthermore, in the three regions of the frontal lobe we recorded from, only a few channels were observed with significant theta-high gamma PAC (59 (6.49%) out of 909 possible channels combined from all frontal areas) and within-condition estimates did not differ as a function of load (pre-SMA: t(4) = -0.16, p = 0.87; dACC: t(13) = -0.82, p = 0.73; vmPFC: t(39) = 0.16, p = 0.87). These observations were qualitatively comparable when significant channels were first averaged within each session, and then tested across recording sessions between the two load conditions (see Fig. S2a), showing that the results were not driven by channels from a single session. There were no significant differences between load conditions in any of the regions for theta-low gamma PAC (30-55 Hz; Fig. S2b). We therefore focused on theta-high gamma (70-140 Hz) PAC for the remainder of the paper.

**Figure 2.**
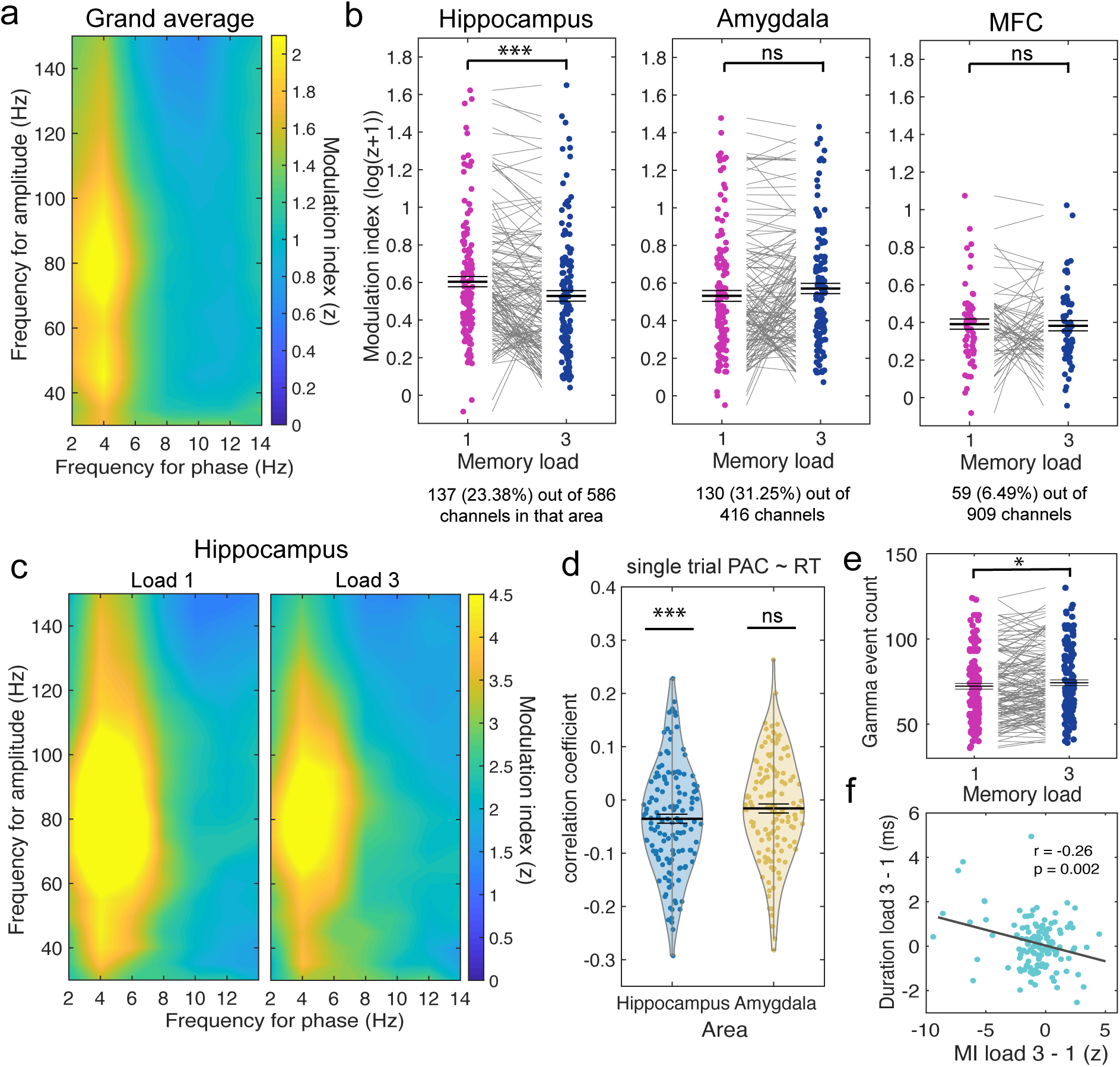
Assessment of theta-gamma phase amplitude coupling. **(a)** Average normalized modulation indices for all phase-amplitude pairs across all n=1949 channels. The strongest phase-amplitude coupling was observed for phases in the theta range (3-7 Hz) and amplitudes from two gamma bands: a low (30-55 Hz) and a high gamma band (70-140 Hz). **(b)** Theta-gamma PAC differed by load. Log-normalized modulation indices were averaged within the theta-high gamma band and compared between the two load conditions in each significant PAC channel (p < 0.05) in each region. PAC channels were common in the hippocampus and amygdala (23% and 31%, respectively) but not in the three frontal areas (6.5%, combined and labeled as MFC (medial frontal cortex)). Only in the hippocampus did theta-high gamma PAC differ significantly as a function of load, with PAC higher in load 1 vs. load 3 (left). Z-scored PAC values were shifted into a positive range by an offset of 1 and log-transformed for illustrative purposes only. All statistics are based on non-transformed z-values. **(c)** Average normalized modulation indices for significant PAC channels from the hippocampus separately for each load condition. **(d)** Theta-gamma PAC was significantly negatively correlated with reaction times in the hippocampus, but not in the amygdala. See Table S1 for GLM results that controls for load differences in RT. **(e)** High-amplitude gamma events were more frequent in load 3 than in load 1 within PAC channels from the hippocampus. **(f)** The longer the gamma events, the weaker was PAC. Shown is the relationship between gamma event duration and PAC. The difference between the durations of high-amplitude gamma events between load 3 and load 1 was negatively correlated with the difference in the modulation index between the two loads across all hippocampal PAC channels. *** p < 0.001; * p < 0.05; ns = not significant, permutation-based t-tests; mixed-model GLMs.

Estimates of PAC can be influenced by nuisance factors such as power or waveform shape differences between conditions (Kramer et al. 2008; Aru et al. 2015; Cole and Voytek 2017). In addition to normalizing PAC estimates within each condition using trial-shuffled surrogates, which controls for confounds such as power differences across conditions, we also directly tested for differences in those nuisance factors in significant PAC channels from the hippocampus (where we observed a significant difference in PAC as a function of load). Power did not differ significantly between the two load conditions in both the theta and the gamma band (all p > 0.05; Fig. S2c). Also, there were no significant differences in theta-gamma phase-phase coupling or theta waveform shape asymmetries between the two load conditions (see Fig. S2d, e). These control analyses confirm that the differences between theta-gamma PAC strength we found were not confounded by power or waveform shape differences.

We next asked whether theta-gamma PAC at the LFP level in the MTL relates to WM behavior. We calculated single-trial estimates of theta-gamma PAC for all significant PAC channels of both MTL regions and then used mixed model GLMs to assess whether reaction time (RT) is related to PAC in a trial-by-trial manner (using only correct trials). To control for potential load differences, we included load as a confounder, and modelled random intercepts for each significant PAC channel. In the hippocampus, PAC was significantly negatively correlated with RT, i.e., faster RTs were associated with stronger PAC (see table S1 for GLM results; Fig. 2d shows correlation coefficients between PAC and RT for each significant PAC channel per region for illustration only; all statistics and conclusions are based on the GLM results). There were no significant correlations between single-trial theta-gamma PAC and RT in the amygdala (see table S1; Fig. 2d).

The above comparison revealed that theta-gamma PAC in the hippocampus was significantly weaker in load 3 as compared to load 1. One possible explanation for this decrease in PAC is that higher WM loads are associated with longer gamma events so that more information per theta cycle can be maintained in WM (Lisman and Jensen 2013). This, in turn, would lead to gamma amplitudes that are more uniformly distributed across the theta cycle and hence weaker levels of PAC (see also Heusser et al. 2016). We next tested this hypothesis in our data. We determined the duration of high-amplitude gamma events within each load condition in the hippocampal PAC channels that showed significant differences in PAC. In correct trials, load 3 trials contained significantly more gamma events than load 1 trials (t(136) = 2.4587, p = 0.01; Fig. 2e). To determine if longer durations of gamma were associated with lower levels of PAC, we next correlated the difference in PAC between the load conditions with the difference in gamma event duration separately for every channel. These two metrics were significantly negatively correlated (r = -0.26; p = 0.002; Fig. 2f). Consistent with our hypothesis, this result suggests that PAC was lower in load 3 trials because the duration of gamma events increased as a function of WM load.

In summary, we observed strong theta-gamma PAC in both hippocampus and amygdala. In the frontal lobe, in contrast, theta-gamma PAC was not prominent. Even more anatomically specific were PAC differences as a function of load and RT, which were significant only in the hippocampus. This finding suggests that PAC relates to ongoing WM processes during the maintenance period in the hippocampus, but not in the amygdala or frontal lobe. We further observed that longer gamma events were associated with weaker PAC estimates in load 3 as compared to load 1, which suggests that more WM information in higher loads was stored across a wider range of theta phases.

### Category neurons synchronize to the phase of gamma signals in the hippocampus

We next sought to determine whether and how spiking activity of simultaneously recorded neurons relates to the LFP phenomena of PAC. Earlier work in humans found that memoranda-specific cells remain persistently active during the WM maintenance period when their preferred stimulus was actively held in WM (Kamiński et al. 2017, 2020; Kornblith et al. 2017). It is believed that these cells are involved in the maintenance of information in WM (Kamiński and Rutishauser 2020). We reasoned that if PAC facilitates interactions between the maintenance of WM content and cognitive control processes, memoranda-tuned neurons that show persistent activity during the maintenance period might be cells that are also related to theta-gamma PAC. Our first approach was hence to determine whether neurons that are tuned to one of the five presented picture categories in the MTL show persistent activity during the maintenance period, and, if so, how they relate to ongoing theta and gamma signals.

We first selected for neurons whose firing rate was related to the visual category of the stimuli shown on the screen during the encoding periods (5-way ANOVA and post-hoc t-test; both p < 0.05; 2,000 permutations). This way of selection leaves the firing rates during the maintenance period independent for later analyses. 89 (24.72 %) out of the 360 neurons in the hippocampus and 181 (36.49 %) out of the 496 neurons in the amygdala were selected as category neurons (see Fig. 3a for an example neuron from the hippocampus). Firing rates of category neurons from both areas were elevated during the maintenance period as compared to baseline across all correct trials (t(269) = 6.65, p < 0.001). Crucially, firing rates were significantly higher in trials in which the preferred category of a cell was held in mind relative to the non-preferred categories (t(269) = 2.93, p = 0.001; Fig. 3b). Category neurons in the MTL thus remained persistently active for their preferred sensory input, even after the sensory material was removed from the screen (areas were combined for simplicity since results were qualitatively similar in hippocampus and amygdala; see Fig. S3a for statistics per area). We further observed a load effect for trials in which the preferred category was encoded, with FRs higher in load 1 than in load 3 (t(269) = 2.65, p = 0.004; Fig. 3c), but not for non-preferred trials (t(269) = -1.46, p = 0.14; Fig. S3b). Overall, FRs were higher in correct as compared to incorrect trials across both load conditions (Fig. 3d; t(245) = 2.4305, p = 0.0185; 24 neurons were excluded from this comparison due to insufficient data in the incorrect condition), further demonstrating their relevance to WM maintenance processes.

**Figure 3.**
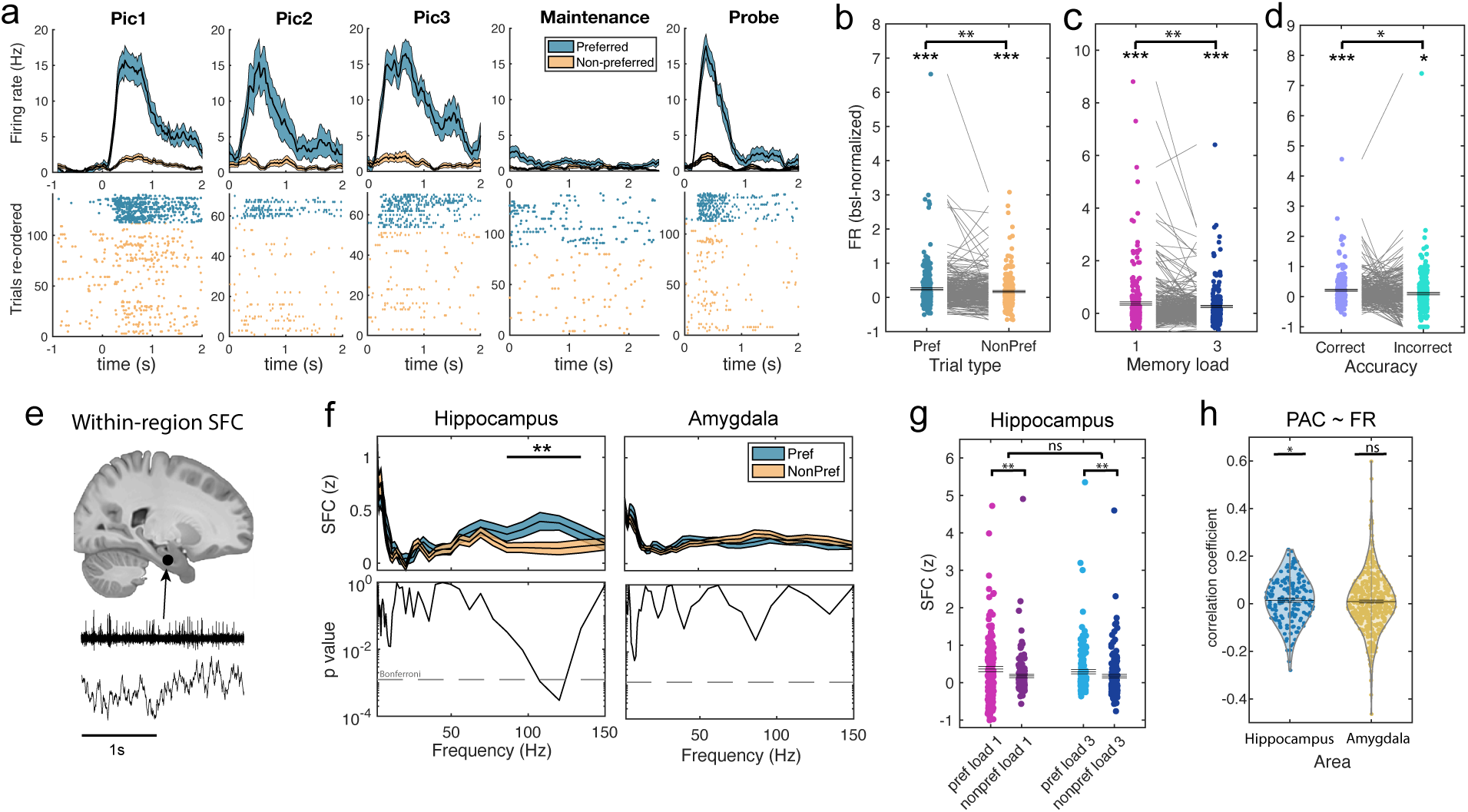
Firing rates and local SFC of category neurons in the MTL. **(a)** Example category neuron recorded in the hippocampus. Category neurons were selected based on higher firing rates for one category than for all other categories during encoding period 1 (ANOVA + post-hoc t-test, both p < 0.05). The preferred category of this neuron was ‘animals’. **(b)** Firing rates averaged across the maintenance period separately for preferred and non-preferred categories for all category neurons from hippocampus and amygdala. Category neurons remained active as compared to baseline and retained their selectivity during the maintenance period of the task, with FR persistently higher for preferred than non-preferred categories. Each dot is a neuron (n=270). Firing rates are shown as percent change to baseline (-0.9 to -0.3 s prior to the first picture onset). **(c)** Firing rates of category neurons during the maintenance period were higher in load 1 as compared to load 3 when their preferred category was held in WM. Each dot is a neuron (n=270). **(d)** In correct trials, FR of category neurons was higher as compared to incorrect trials across both load conditions. Each dot is a neuron (n=246). 24 neurons were excluded due to insufficient data in the incorrect condition. **(e)** We computed local spikefield coherence between spikes and LFPs recorded in the same area for all category neurons and compared preferred versus non-preferred trials. **(f)** When paired with local PAC channels, spikes of category neurons in the hippocampus were significantly more strongly phase-locked to local gamma LFPs in the high gamma range during the maintenance period when the preferred category of a neuron was held in WM. No significant differences were found for the amygdala or non-PAC channels (see Fig. S3e). **(g)** Gamma (70-140 Hz) SFC for hippocampal category neurons was significantly stronger for preferred vs non-preferred trials in both load conditions. No main effect of load or interaction was found. Each dot is a neuron-LFP channel pair (n=151). **(h)** Theta-gamma PAC was significantly positively correlated with firing rates of category neurons in the hippocampus, but not in the amygdala. See Table S2 for GLM results that control for load differences in PAC. *** p < 0.001; ** p < 0.01; * p < 0.05; ns = not significant; (cluster-based) permutation t-test/ANOVA; mixed-model GLMs.

Category cells thus showed persistent activity during the maintenance period of the task. Next, we asked how spike timing of category neurons relates to the phase of LFPs recorded within each area of the MTL (Fig. 3e). We computed spike-field coherence (SFC) for all category neuron to channel combinations within the same region in frequencies between 2 and 150 Hz during the maintenance period and compared trials in which preferred or non-preferred stimuli were correctly maintained. Cluster-based permutation statistics revealed that high gamma-band SFC was significantly stronger in preferred trials than non-preferred trials across all neuron-to-channel combinations that involved significant PAC channels in the hippocampus (cluster-p = 0.004; Fig. 3f). This data-driven approach thus revealed a gamma cluster that spanned roughly the same frequencies involved in theta-high gamma PAC (70-140 Hz vs 86-134 Hz). We did not find similar effects for the theta band (Fig. 3f), non-PAC channels (Fig. S3e), or neuron-channel combinations from the amygdala (Fig. 3f; S3e). To determine whether the observed gamma SFC difference between preferred and non-preferred trials was dependent on gamma amplitude, we tested whether gamma SFC (averaged across 70-140 Hz) for category neurons in the hippocampus differed between preferred and non-preferred trials for high and low gamma amplitudes separately (median split). We observed a significant difference only for spikes that occurred during high gamma power (t(150) = 3.0601, p = 0.0015), not low power (t(150) = 0.26, p = 0.85; Fig. S3d). Therefore, specifically in periods where gamma amplitude was high, spikes of category neurons were more strongly synchronized to the phase of gamma signals when their preferred as compared to non-preferred category was maintained in WM.

We next tested whether gamma SFC also differed as a function of WM load. We averaged SFC within the gamma band (70-140 Hz) and computed a 2×2 ANOVA with the factors *load* and *preference* for all category neuron to PAC channel combinations in the hippocampus. We observed a significant main effect for preference (F(1,150) = 16.23, p < 0.001), whereas there was no significant main effect of load (F(1,150) = 1.32, p = 0.25) nor a significant interaction between the two (F(1,150) = 0.93, p = 0.33). Confirming this result, gamma-band SFC was significantly elevated for preferred vs. non-preferred trials in load 1 (t(150) = 3.14, p = 0.003) as well as in load 3 (t(150) = 2.88, p = 0.004; Fig. 3g). Computing the same 2×2 ANOVA for SFC values averaged within the theta band (3-7 Hz) did not reveal any significant effects (Fig. S3c).

Lastly, we determined whether spiking activity of category cells during the WM maintenance period correlates with PAC on a trial-by-trial manner in both areas of the MTL (only using correct trials). We again included load as confounder and modelled random intercepts for each category neuron to significant PAC channel combination. In the hippocampus, PAC was weakly but significantly positively correlated with FR of category neurons (see table S2 for GLM results; Fig. 3h shows correlation coefficients for illustration only; all conclusions are based on the GLM results). In the amygdala, there were no significant correlations between single-trial theta-gamma PAC and FR of category neurons (see table S2; Fig. 3h). Together, these results show that category-selective neurons in the hippocampus were persistently active and more synchronized with gamma LFPs when their preferred category was held in WM. This effect was specific to channels that showed significant theta-gamma PAC during the maintenance period (Fig. S3e). FRs of hippocampal category neurons were moreover correlated with single-trial estimates PAC.

### Category neurons did not significantly overlap with PAC neurons

While above result indicates a relationship between category neurons and PAC within the hippocampus, these results alone do not definitively demonstrate that spiking activity of category neurons was sensitive to the nonlinear interaction between theta phase and gamma amplitude as would be expected from a true “PAC neuron”.

We thus next selected for PAC neurons, which we defined as neurons whose firing rate was sensitive to both theta phase and gamma amplitude. We then examined whether the selected neurons were significantly overlapping with the population of category neurons. To select for PAC neurons, we fit three different models (Poisson GLMs) to each neuron-to-channel combination within a region and session. We extracted the theta phase and gamma amplitude for each spike of a given neuron during the maintenance period of correct trials in each corresponding channel and used them as predictors for the spike count of the corresponding neuron binned into 20 bins (see Methods). We reasoned that if a neuron’s spiking activity is correlated with theta-gamma PAC, the interaction term between theta phase and gamma amplitude should explain significant amounts of variance in the spike count of a given neuron. This is because the activity of a PAC neuron should not only be related to theta phase or the gamma amplitude alone (explained by the main effects of the model), but specifically also to the interaction of the two, i.e., a gamma amplitude that is differentially distributed across theta phase, hence PAC. To test this hypothesis, we performed a model comparison (likelihood-ratio test) of the full model that included both the two main effects theta phase and gamma amplitude as well as their interaction against a model that included the two main effects but no interaction term. We also tested the full model against a model that did not include the main term for gamma but only the main term for theta and their interaction (see Methods for the reasoning behind this comparison, Fig. S4 for illustrations, and Fig. 4a for an example). We selected a neuron as a PAC neuron if the full model explained the spike count variance significantly better than both other models for at least one of the channel combinations (p < 0.01; FDR corrected for all possible channel combinations; if a neuron had more than one significant LFP-spike channel combination, we selected the one with the highest R^2^ in the full model for later within-region SFC analyses). A substantial proportion of neurons in the MTL qualified as PAC neurons: In the hippocampus, 79 (37.29 %) out of 212 available neurons (p < 0.005; 200 permutations, see Methods; pre-processing steps removed broadband LFPs for some of the neurons and those were therefore not part of this analysis) qualified as PAC neurons. In the amygdala, 163 (45.53 %) out of 358 neurons (p < 0.005) qualified as PAC neurons.

**Figure 4.**
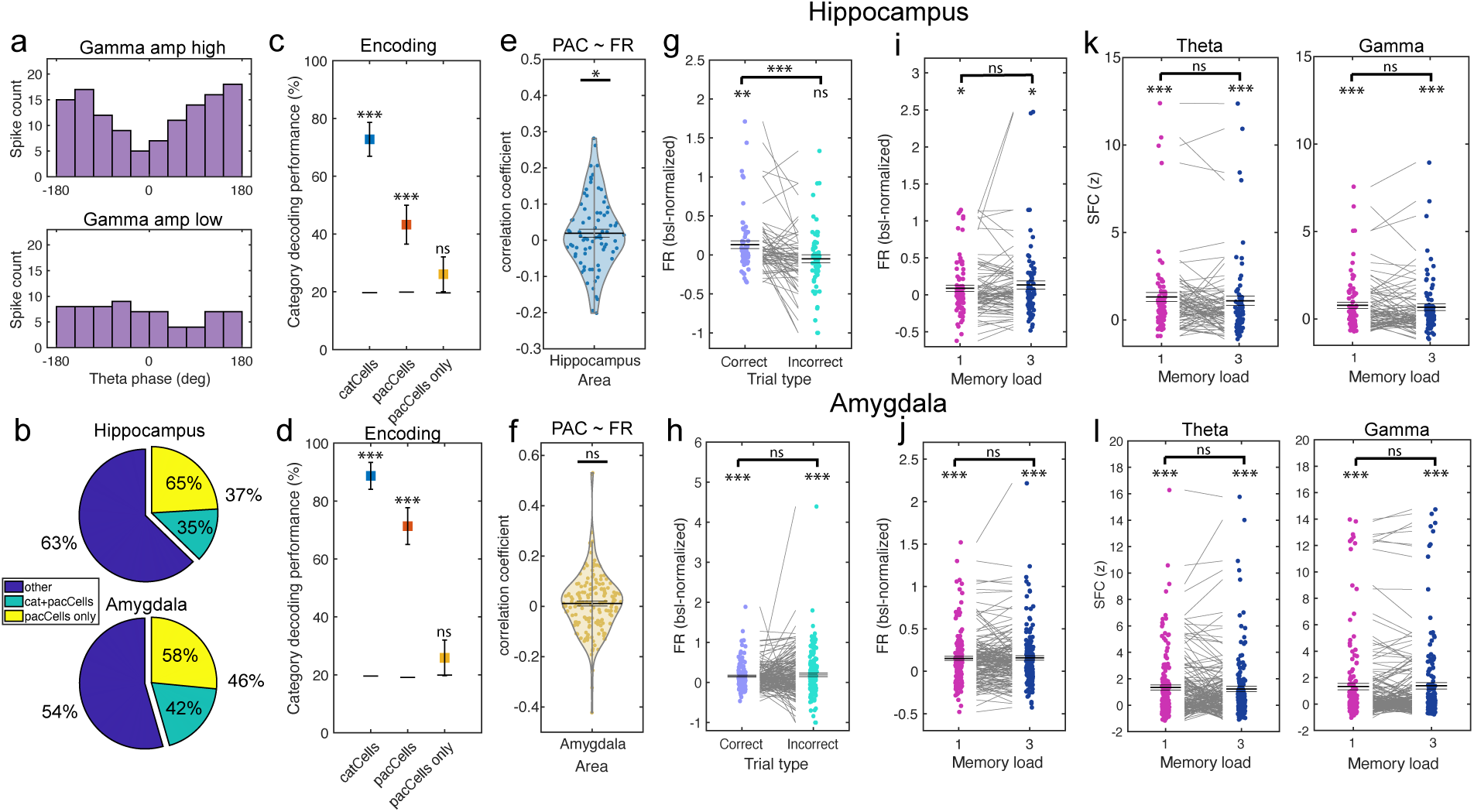
PAC neuron selection and local activity. **(a)** Example showing binning used for PAC neuron selection for a neuron from the hippocampus. Theta phase, binned into ten groups, and gamma amplitude, median split into low and high, were used to predict spike counts of each neuron from the MTL during the maintenance period. Only if the model containing the two factors and their interaction predicted spike counts significantly better than two other models that lacked the interaction or the gamma amplitude main effect term, respectively, a given neuron was selected as PAC neuron. In this example neuron from the hippocampus, spike count was higher during high gamma amplitudes (gamma main effect) and differed in their theta phase distribution between high and low gamma amplitudes (interaction effect), resulting in selecting this neuron as a PAC neuron. In the analysis, we separated the theta phase into sine and cosine terms to account for the circularity of phase values, which is not shown here for simplicity. **(b)** Proportions of all recorded neurons that qualified as PAC neurons (yellow, green). **(c,d)** PAC neurons were not selective for category. Even during picture presentation (encoding), image category could not be efficiently decoded from firing rates of “PAC only” neurons. Error bars reflect the standard deviation of 1,000 decoding repetitions. Black horizontal lines indicate mean decoding accuracy of 1,000 randomly shuffled category labels. Decoding was performed for pseudo-populations of category or PAC neurons, respectively. **(e,f)** Firing rates of PAC neurons were positively correlated with single-trial estimates of theta-gamma PAC in the hippocampus, not the amygdala (see Table S3 for GLM results). **(g,h)** Firing rates of PAC neurons during the maintenance period differed between correct and incorrect trials in the hippocampus but not amygdala. Firing rates are shown as percent change to baseline (-0.9 to -0.3 s prior to the first picture onset). **(i-l)** Firing rates as well as theta, gamma SFC between PAC neurons and local LFP recordings did not differ as a function of load in both MTL areas. Theta and gamma SFC, however, were both significantly stronger than shuffled surrogates in the hippocampus as well as the amygdala. In (e-l), each dot is a neuron. *** p < 0.001; ** p < 0.01; * p < 0.05; ns = not significant; permutation-based t-tests; mixed-model GLMs.

Next, we determined whether the selected PAC neurons were also category neurons. In the hippocampus, 28 (35.44 %) out of the 79 PAC neurons were both PAC neurons as well as category neurons. In the amygdala, this was the case for 68 (41.72 %) out of the 163 PAC neurons (Fig. 4b). We compared these proportions to 10,000 random selections of the same number of neurons from all available neurons in each region and found that the proportion of category neurons among PAC neurons was not significantly higher than expected by independent sub-populations in any of the two regions (both p > 0.05). This suggested that the probabilities of a neuron being a PAC or a category neuron were independent, therefore not confirming our initial hypothesis.

To further corroborate this finding, we trained a linear decoder to differentiate between the five different picture categories based on the firing rates extracted during picture presentation (encoding). While, as expected, the decoder was able to differentiate between the picture categories when trained on FR from the category neurons with high accuracy (hippocampus: 72.77 %; p = 0.001; amygdala: 88.71 %, p = 0.001; significance assessed by comparing original decoding accuracy to a distribution of 1,000 decoding accuracies after randomly shuffling category labels; Fig. 4c, d), it could not differentiate the categories when trained on FRs from PAC neurons that were not also category neurons in both MTL areas (hippocampus: 26.06 %, p = 0.16; amygdala: 25.86 %, p = 0.15; chance level = 20 %). Thus, PAC neurons were not differentially active for the five picture categories and therefore differed in their response from category neurons.

### Properties of PAC neurons

We next sought to characterize PAC neuron activity and its relation to WM maintenance processes within the MTL during the maintenance period. We first asked whether the FR of PAC neurons correlates with the LFP-based estimates of theta-gamma PAC on a trial-by-trial manner. Trial-by-trial correlations are independent from the selection procedure since PAC neurons were selected based on trial-averaged theta-gamma interactions, irrespective of their trial-by-trial variance. For this purpose, we again computed a mixed-model GLM, including load as a confounder and modelling a random intercept for each PAC neuron-to-channel combination (using only correct trials and the LFP channel selected for each neuron; see Methods). We found that FRs of PAC neurons in the hippocampus were positively correlated with estimates of single-trial PAC (see table S3 for GLM results; Fig. 4e,f shows univariate correlation coefficients for illustration only). In terms of FR, PAC neurons in the hippocampus showed elevated activity throughout the maintenance period as compared to baseline (t(78) = 2.43, p = 0.01), had significantly higher FRs for correct as compared to incorrect trials (t(62) = 3.82, p < 0.001; Fig. 4g; 16 neurons were excluded from this comparison due to insufficient data in the incorrect condition), and were elevated as compared to baseline in correct (t(62) = 2.67, p = 0.01) but not in incorrect trials (t(62) = -0.98, p = 0.33). FRs were not significantly different between the two load conditions (load 3 - load 1: t(78) = 1.38, p = 0.20), but FRs were elevated as compared to baseline in each condition considered separately (load 1: t(78) = 2.14, p = 0.03; load 3: t(78) = 2.45, p = 0.01; Fig. 4i). In the amygdala, FRs of PAC neurons were not significantly correlated with single-trial estimates of theta-gamma PAC (Table 3; Fig. 4f). PAC neurons in the amygdala also showed higher FRs during the maintenance period as compared to baseline (t(162) = 6.40, p < 0.001), but we did not observe significant differences between correct and incorrect WM trials (t(156) = -0.77, p = 0.45; Fig. 4h) or loads (load 3 - load 1: t(162) = 0.26, p = 0.80: Fig. 4j). These results show that PAC neurons in the hippocampus, despite not being tuned to WM content, were actively contributing to WM maintenance because their FRs were elevated during the WM maintenance period as compared to baseline and higher in correct as compared to incorrect trials.

We further asked whether the SFC of PAC neurons differed with respect to local theta or gamma between the two load conditions. We computed theta and gamma SFC for all neuron-to-channel combinations determined during the selection process for PAC neurons. While SFC was significantly stronger as compared to shuffled surrogate data in each load condition in the hippocampus in the theta (load 1: t(78) = 5.1505, p < 0.001, load 3: t(78) = 4.4853, p < 0.001; Fig. 4k) and the gamma band (load 1: t(78) = 4.4915, p < 0.001; load 3: t(78) = 3.5826, p < 0.001), we did not observe significant differences between the two load conditions (load 3 – load 1: theta: t(78) = -1.54, p = 0.13; gamma: t(78) = -1.12, p = 0.27). PAC neurons in the amygdala showed comparable results (comparisons within load conditions: all p < 0.001; load 3 – load 1: theta: t(162) = -0.71, p = 0.47; gamma: t(162) = 0.76, p = 0.45: Fig. 4l).

### PAC neuron activity in the hippocampus is related to frontal theta LFPs

Above analysis shows that unlike category neurons, the firing rate of PAC neurons in the MTL does not seem to be related to the content of WM. However, the activity of PAC neurons was predictive of WM quality because it differed between correct and incorrect trials in the hippocampus. We therefore hypothesized that PAC neurons might be involved in cognitive control of WM maintenance processes. While it has long been theorized that long-range theta connectivity might be the substrate underlying this control process (Miller 2000; Fell and Axmacher 2011; Daume et al. 2017b, a), no single cell correlate of this mechanism is known. We thus asked whether the activity of PAC neurons is related to activity recorded in frontal regions (Minxha et al. 2020).

We computed cross-regional SFC between spiking activity of PAC neurons in the MTL and the phase of LFPs between 2 and 150 Hz recorded in the three frontal regions pre-SMA, dACC and vmPFC (Fig. 5a). We then tested for differences in cross-regional SFC between the two WM load conditions. If PAC neuron activity is related to frontal cognitive control, we expected cross-regional SFC in the theta range to be stronger in load 3 than in load 1 since higher levels of cognitive control are required for higher WM loads. The data supports this hypothesis: SFC was significantly stronger in load 3 than in load 1 between spiking activity of PAC neurons in the hippocampus and theta-band LFPs recorded in the vmPFC (cluster-based permutation statistics, cluster-p < 0.001; Fig. 5b; for analysis separate for narrow- and broad-spiking neurons, see Fig. S5). We did not observe significant differences for other frequency bands, nor for the other two frontal brain areas (see Fig. S5a). To determine if this effect was specific to PAC neurons, we repeated the same analysis for the category neurons and did not observe any significant differences (Fig. 5c). Similarly, PAC neurons from the amygdala did not show significant cross-regional SFC differences in any of the tested frequencies (Fig. 5d) or regions.

**Figure 5.**
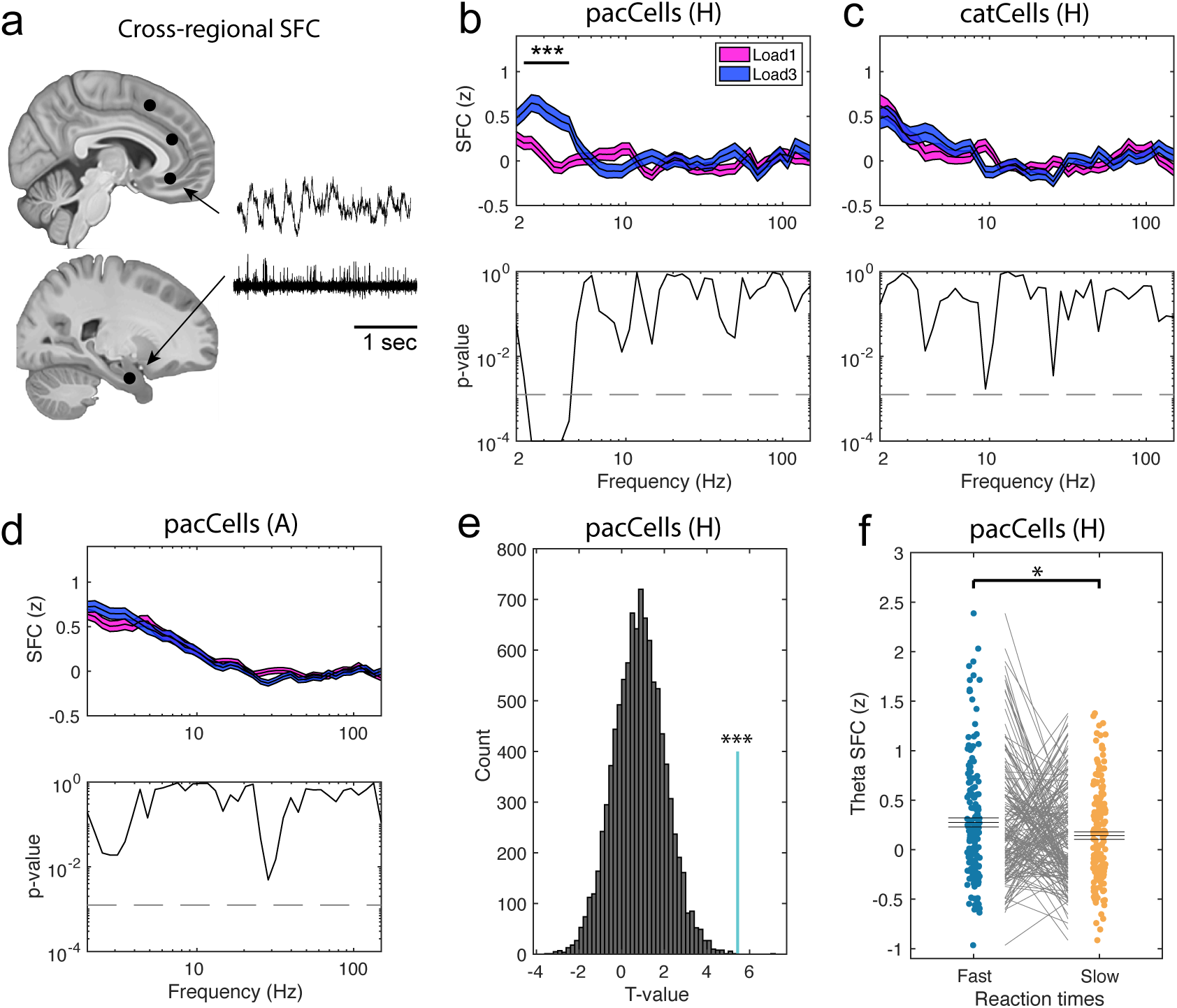
Remote connectivity of PAC neurons in the MTL to frontal theta LFPs. **(a)** We computed long-range SFC between MTL spiking activity and LFPs recorded from all three frontal regions. **(b)** Spikes of PAC neurons in the hippocampus were more strongly synchronized with theta-band LFPs recorded in the vmPFC during the maintenance period during load 3 compared to load 1 trials. This was not the case for pre-SMA and dACC (see Fig. S5a). **(c)** Category neurons from the hippocampus, or **(d)** PAC neurons from the amygdala did not show significant SFC differences between loads relative to vmPFC LFP. **(e)** Hippocampal PAC cells (n=175 connections, cyan line) yielded the strongest long-range theta SFC difference between load 3 and load 1 trials among 10,000 random selections of hippocampus-vmPFC connections. T-values correspond to comparisons between load 3 and 1 for an average of SFC values in the significant theta range (2.5-4.3 Hz). **(f)** Remote theta-band SFC between spiking activity of PAC neurons and LFPs recorded in the vmPFC was significantly stronger for fast than for slow RT trials. Each dot is a neuron-channel connection (n=167; some neuron-channel pairs were excluded due to inefficient spike count in at least one of the conditions). *** p < 0.001; * p < 0.05, H = Hippocampus; A = Amygdala; (cluster-based) permutation t-tests.

To further corroborate whether the observed difference in cross-regional theta SFC was specific to PAC neurons, we compared the strength of cross-regional SFC between the load conditions for SFC values averaged in the observed theta range for 10,000 random selections of hippocampal neurons (same number of connections as for PAC neurons). None of the random selections yielded a stronger difference between the two load conditions than the PAC neurons from the hippocampus (p < 0.001; Fig. 5e).

If cross-regional SFC reflects levels of cognitive control during WM maintenance, theta SFC should not only be stronger for higher loads but also for faster RTs. Thus, we asked whether cross-regional theta SFC between PAC neurons and LFPs in the vmPFC differed between fast and slow RTs. We performed a median split of RTs for all correct trials within each load condition and compared cross-regional SFC for PAC neurons and theta in the vmPFC between fast and slow RTs (averaged across both load conditions). Theta SFC was stronger for fast as compared to slow RTs for PAC neurons from the hippocampus (t(166) = 2.10, p = 0.03; Fig. 5f), but not from the amygdala (t(705) = 1.40, p = 0.16).

### PAC neurons shape representations of WM content and facilitate WM behavior through noise correlations

Above results show that PAC neurons are involved in cognitive control by orchestrating long-range fronto-temporal interactions. However, this leaves open the question of how this PAC-neuron mediated control process might shape WM representations at the local level, thereby facilitating WM maintenance. We hypothesized that PAC neurons could serve to stabilize WM content representations by introducing information-enhancing noise correlations. Noise correlations among a group of simultaneously recorded neurons can significantly modify the information content of the population because correlations can shape representations such that they can become easier or harder to read out (Averbeck et al. 2006; Panzeri et al. 2022). Notably, even cells that by themselves carry no information in their firing rate (they are untuned) can influence the decodability of variables at the population level if their activity is correlated trial-by-trial with cells that are tuned (Leavitt et al. 2017b; Stefanini et al. 2020). We hypothesized that PAC neurons might play this role during WM maintenance, during which cognitive control is needed to hold information in memory.

We first tested whether pairs of PAC and category cells that were recorded in the same session and brain area had significantly correlated firing rates. In each trial and neuron, we counted spikes in windows of 200 ms, sliding across the maintenance period in steps of 25 ms. We then correlated spike counts for pairs of neurons across all time bins and averaged correlation coefficients for all correct trials. We excluded pairs of neurons that were recorded on the same channel to avoid potentially spurious correlations due to misclustered spikes. In both the hippocampus (162 pairs; t(161) = 5.26; p < 0.001) and the amygdala (892 pairs; t(891) = 15.51; p < 0.001), we found significant on average positive single-trial co-fluctuations of spike counts among pairs of category neurons and PAC neurons (Fig. 6a). To further corroborate this finding, we shuffled trial labels within each pair and recomputed the GLMs for 1,000 times. The correlation coefficients averaged across all PAC and category neuron pairs with intact trial labels was significantly stronger than for shuffled trials (see Fig. 6b for hippocampal pairs; for noise correlations computed across trials, see Fig. S6).

**Figure 6.**
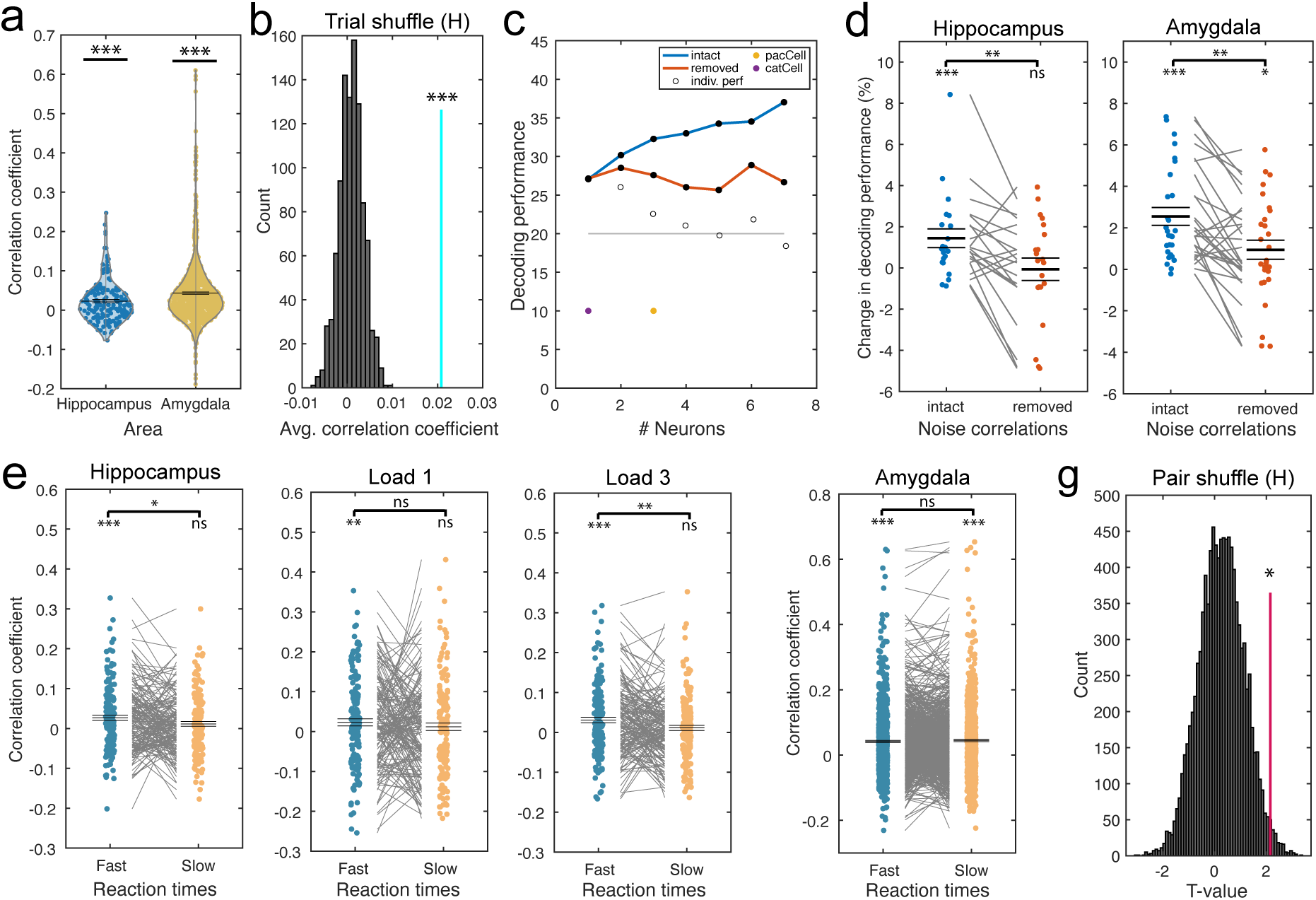
Noise correlations of PAC neurons within MTL. **(a)** Trial-averaged, bin-wise correlation coefficients for all possible pairs of category and PAC neurons in the hippocampus and amygdala. In both regions, correlation coefficients were significantly higher than zero on average. **(d)** Repeat of the correlation analysis for all possible PAC-category neuron pairs in the hippocampus. Shuffling trial labels for 1,000 times resulted in far lower correlations between pairs of neurons than unshuffled trial labels (cyan line; mean of correlation coefficients across all pairs. **(c)** Single-session example for optimized decoding accuracies for firing rates during the maintenance period for hippocampal neurons. Category decoding accuracies were computed with intact or removed noise correlations among neurons. Purple dots indicate when a category cell, yellow dots when a PAC neuron was added to the optimized decoding ensemble. White dots show decoding accuracies for each individual neuron. **(d)** (left side) Adding hippocampal PAC neurons to the optimized decoding ensemble significantly enhanced decoding performance of WM content when noise correlations were kept intact. When noise correlations were removed, in contrast, PAC cells did not improve decoding performance. This indicates that, for PAC cells, noise correlations rather than individual firing rates shaped the geometry of category representations. (right side) Amygdala PAC neurons contributed more to decoding memory content with intact noise correlations. Note, however, that amygdala PAC cells contributed to decoding also after removing noise correlations during the maintenance period. Each dot represents a neuron. **(e)** Noise correlations among hippocampal PAC and category neurons were stronger in fast than slow RT trials (median split) for trials in which the category neuron’s preferred category was correctly maintained (left). This effect was especially strong in load 3 trials (right), not in load 1 trials (middle). **(f)** In the amygdala, we did not observe a significant difference in noise correlations between fast and slow trials when the preferred category was maintained for pairs of PAC and category neurons. In (e,f) each dot represents a cell pair. **(g)** Pairs of PAC and category neurons in the hippocampus (pink line) showed a significantly stronger difference in noise correlations between fast and slow RT trials than randomly selected pairs of any recorded neuron (from the same session) and category neurons in 10,000 repetitions. *** p < 0.001; ** p < 0.01; * p < 0.05; ns = not significant; H = Hippocampus; permutation-based t-tests.

As a next step, we sought to determine whether PAC cells contribute to the decodability of image category during the WM maintenance period through structuring noise correlations. We used the approach introduced by (Leavitt et al. 2017b), which consists of iteratively adding neurons to the population through greedy selection of the neuron that adds most decodability above and beyond that provided by the already included neurons. First, the unit that is most informative for stimulus category is determined by decoding the category from each recorded neuron by itself. Second, all possible pairs of this best neuron and all remaining neurons are individually used for decoding, picking the best second neuron. Third, this procedure is repeated iteratively till all neurons are added to the population (see Fig. 6c for an example from a single session). Previous work using this method has observed that neurons that by themselves provide no information are added to the population (Leavitt et al. 2017b; Stefanini et al. 2020). We performed this analysis with intact noise correlations among neurons (by using original trial labels) and after noise correlations were removed (by shuffling trials within each category to preserve correct decoding labels but destroy correlations across trials).

We determined each PAC neuron’s contribution to category decodability after it was added to the optimized decoding ensemble (difference to decoding accuracy before that PAC neuron was added) and compared their contributions when noise correlations were intact or removed within each brain area (for all neurons added before peak decodability; see (Leavitt et al. 2017b)). In the hippocampus, adding PAC neurons to the optimized decoding ensemble significantly enhanced category decoding during the WM maintenance period when noise correlations were intact (t(20) = 3.16, p < 0.001) but not when they were removed (t(20) = -0.12, p = 0.91; Intact vs. Removed: t(20) = 3.33, p = 0.003; Fig. 6d left). In the amygdala, PAC neurons contributed to category decodability not only when noise correlations were intact (t(27) = 5.90, p < 0.001) but also when noise correlations were removed (t(27) = 2.04, p = 0.04; Intact vs Removed: t(27) = 4.31, p = 0.0011; Fig. 6d right). These results show that PAC neurons in the hippocampus contributed to the encoded WM content only through noise correlations to tuned cells during the WM maintenance period. When noise correlations were removed, these contributions were abolished.

We next sought to determine whether noise correlations among PAC and category neurons benefited WM-based behavior. If noise correlations are beneficial to WM processes, they should be stronger in correct fast RT trials as compared to correct slow trials, specifically when the category neurons’ preferred categories were maintained in WM. We thus compared noise correlations between fast and slow RT trials (median split; separately computed in each load condition and then averaged to avoid a bias of load in RTs) for the category neurons’ preferred trials. In the hippocampus, we observed significantly stronger noise correlations for fast as compared to slow RT trials (t(161) = 2.15; p = 0.028; Fig. 6e left). Noise correlations were on average significantly positive only in fast (t(161) = 4.10, p < 0.001;), but not in slow trials (t(161) = 1.94, p = 0.06). For non-preferred trials, we did not observe a significant difference between fast and slow RT trials (see Fig. S6d). Separating trials into the two load conditions, we only observed a significant difference between fast and slow trials in load 3 (t(161) = 2.60, p = 0.009; Fig. 6e right), not load 1 (t(161) = 0.96, p = 0.34; Fig. 6e middle). In the amygdala, comparing fast to slow RT trials in preferred trials did not reveal a significant difference (t(891) = -1.00, p = 0.33; Fig. 6f).

Lastly, we asked whether the effect of noise correlations on RTs in the hippocampus was specific to PAC-to-category neuron pairs or a common feature across the entire population of simultaneously recorded neurons. For this purpose, we randomly paired category neurons with any other non-PAC neuron recorded in the hippocampus within the same session and compared noise correlations between fast and slow RT trials (for n = 162 randomly selected pairs; same n as for PAC-to-category neuron pairs). Repeating this analysis for 10,000 times, we observed that PAC-to-category neuron pairs showed a significantly stronger effect than most randomly selected cell pairs (p = 0.016; Fig. 6g). This result shows that, in particular, the noise correlations between category and PAC neurons within the hippocampus contributed to enhanced WM fidelity.

## Discussion

Our data reveal a PAC-mediated mechanism for the control of WM maintenance. We identified “PAC neurons”, whose spiking activity followed the interactions between theta phase and gamma amplitude during the maintenance period of a Sternberg WM task. Unlike category neurons, which displayed memoranda-specific persistent activity, the activity of PAC neurons was not related to WM content per se. Rather, the activity of PAC neurons in the hippocampus was related to the cognitive control processes that enable the efficient and accurate maintenance of WM. Firing patterns of hippocampal PAC neurons were more strongly synchronized with the phase of frontal theta signals in trials with higher load and faster RT. Moreover, trial-by-trial co-fluctuation in FR between PAC and category neurons in the hippocampus shaped the population-level geometry of WM representations such that WM fidelity was improved, with stronger pair-wise noise correlations resulting in faster RTs. We conclude that these interactions between PAC and category neurons are a reflection of the interplay between top-down control and the local processing of WM content that make WM maintenance possible.

In line with earlier studies (Tort et al. 2008; Cabral et al. 2014; Yamamoto et al. 2014), activity in the high gamma (70-140 Hz) frequency range was reflective of processing and WM maintenance of sensory information. Yamamoto and colleagues (Yamamoto et al. 2014) found that in mice, the activity of high gamma oscillations (65-140 Hz) in the hippocampal-entorhinal system was related to the successful execution of WM maintenance. Synchronization in the high gamma band between the entorhinal cortex and hippocampus was stronger in correct than incorrect trials, and appeared shortly before a reversal of a decision when the animal initially made a wrong choice. The authors suggested that high gamma activity thus contributes to the explicit awareness of WM content. They did not find a relation of WM processes to activity in the lower gamma band (25-50 Hz). Similarly, Tort et al. (Tort et al. 2008) reported that PAC between theta and high gamma oscillations in the rat hippocampus, which was especially strong in time periods after a sensory cue has been represented, presumably involving processes of WM maintenance and decision making. They also did not observe PAC between theta and low gamma. Based on those and other results, the high gamma band has thus been suggested to signal the routing of encoded sensory information into and within the hippocampus (Colgin et al. 2009; Colgin 2016; Fernández-Ruiz et al. 2021). In favor of this hypothesis, PAC in our study differed as a function of WM load only for frequencies involving the high gamma range, signaling that the processing of WM content affected neural activity specifically in those frequencies. High-amplitude events in the high gamma range were more abundant in load 3 as compared to load 1 trials, and their difference in duration between the load conditions was negatively correlated with the difference in PAC estimates across channels. In line with earlier reports (Heusser et al. 2016), this suggests that storing more information in WM leads to lower estimates of theta-high gamma PAC since longer high gamma events are more broadly distributed across the theta cycle. Studies in rats observed an increase in the number of gamma cycles specifically in the high gamma range, not the low range, when the length of a running track increased (Gupta et al. 2012; Zheng et al. 2016), potentially signaling an increase in WM load during spatial navigation. Moreover, in our study persistently active category neurons were more strongly phase locked to signals in the high gamma range when their preferred stimulus was maintained. No effects were observed for frequencies involving the low gamma band (30-55 Hz). Our study thus provides evidence for a specific role of the high gamma band in the processing and maintenance of WM content in the hippocampus.

Although we observed significant theta-gamma PAC in the amygdala as well, activity in the high gamma band in the amygdala was not related to WM processes because theta-gamma PAC differed neither as a function of WM load nor was it related to WM-based behavior. Also, unlike in the hippocampus, category cells in the amygdala were not more strongly coupled to gamma when their preferred stimulus was maintained in WM. Earlier reports indicated that the amygdala plays a role in the maintenance of information in WM (Schaefer and Gray 2007; Kamiński et al. 2017). Our observation of stimulus-specific persistent activity of category neurons in the amygdala provides further evidence for this claim. However, our results indicate that the amygdala supports WM through a different mechanism than the hippocampus. Note that the absence of a role of PAC in the amygdala was not due to an absence of PAC cells, which were common in the amygdala. Whether PAC and high gamma in the amygdala serve a different role than that of the hippocampus during WM maintenance remains an open question.

We identified “PAC neurons” in the medial temporal lobe, whose spiking activity was predicted by the interaction between theta phase and gamma amplitude. Up until now it was unclear how observations of theta-gamma PAC at the LFP level translated to single neuron activity or whether those interactions were visible at the single neuron level at all. Here, we now show for the first time that theta-gamma PAC has a direct relation to the spiking activity of individual neurons. While, by definition, these PAC neurons were related to local gamma activity, they were not directly involved in the processing of the current memoranda per se. PAC neurons did not significantly overlap with the population of category cells more than expected by independence and were not preferably active for certain picture categories during stimulus presentation. Their FR was not informative about the current WM memoranda such that a linear classifier failed to decode the currently presented picture category from their spiking activity. Importantly, however, the activity of PAC neurons was coordinated with frontal theta LFPs in the vmPFC. This phase locking was stronger for higher WM load and faster RTs, indicating a role in cognitive control. Relatedly, Liebe and colleagues (Liebe et al. 2012) observed enhanced cross-regional phase coupling in the theta range between single neurons in macaque V4 and LFPs recorded in lateral prefrontal cortex during a WM maintenance period. While this prior study shows that phase locking of V4 neurons to frontal theta activity was stronger in successful as compared to error trials, this study left it unclear whether such phase coupling was related to the cognitive control of WM content. Also, in this prior study, theta coupling was not related to modulations of WM load nor to interactions with local maintenance processes of WM content in higher frequencies. Together, our data show for the first time that hippocampal PAC cells specifically play a role in engaging cross-regional theta phase locking as a way to control the hippocampal processing of WM content.

Long-range theta phase locking between frontal and temporal/occipital areas has been suggested to reflect frontal cognitive control exerted over task-relevant brain processes in widespread sensory and association areas (Miller 2000; Fell and Axmacher 2011; Helfrich and Knight 2016). In WM maintenance, fronto-temporal interactions are crucial, especially when involving the hippocampus (Harris and Gordon 2015), and often involve the theta and gamma frequency range (Sauseng et al. 2004; Liebe et al. 2012; Spellman et al. 2015; Hallock et al. 2016; Daume et al. 2017a, b; Johnson et al. 2017; Tamura et al. 2017; Malik et al. 2022). Theta-based prefrontal coordination of posterior WM content-specific processes could ensure efficient information processing at phases that are optimal for network-wide communication within the memoranda-processing population of neurons (Fell and Axmacher 2011; Fries 2015). According to this model, higher levels of cognitive control are indicated by stronger phase locking between regions to facilitate faster and more efficient readout of WM content. Our results support this model and provide, for the first time, evidence for a specific mechanism to implement it. Spike-field coherence between hippocampal PAC neurons and theta LFPs in the vmPFC was stronger in load 3 than in load 1, i.e., in periods where more cognitive control was required to coordinate a higher load of WM information. We note that while vmPFC is known to be involved in top-down control processes (Badre and Nee 2018; Chai et al. 2018; Yin et al. 2021), especially in interaction with the hippocampus (Gluth et al. 2015; Jin and Maren 2015; Günseli and Aly 2020), we are the first to show that it is engaged in the long-range cognitive control of the maintenance of WM information in the hippocampus. Cross-regional hippocampus-vmPFC SFC was enhanced for fast as compared to slow RT trials, which signals a more efficient read-out of WM content with stronger levels of control. These results thus indicate that prefrontal-hippocampal communication in the theta band reflects cognitive control routed to WM content-processing areas such as the hippocampus during WM maintenance.

One way by which cognitive control is exerted is thought to be via monosynaptic projections from PFC to inhibitory interneurons in the hippocampus (Malik et al. 2022). Malik et al. observed that more top-down control led to enhanced signal to noise ratios of object-related spatial encoding and, at the same time, reduced overall network activity and inhibited feedforward processing in the hippocampus. Relatedly, we observed cognitive control related signals between hippocampal PAC neurons and vmPFC specifically for narrow-spiking neurons (see Fig S5), which are thought to likely reflect inhibitory interneurons (Barthó et al. 2004; Mosher et al. 2020). This therefore suggests the new specific hypothesis that the hippocampal PAC neurons we described are inhibitory interneurons that receive monosynaptic projections from PFC. Further supporting this hypothesis is our finding that category neurons reduce their firing rate in load 3 compared to load 1 trials. It is possible that this is due to increased inhibition exerted as a function of increased need for cognitive control.

MEG studies of WM maintenance indicate that in the human temporal lobe, local PAC co-exists together with long-range theta phase synchronization to the frontal lobe (Daume et al. 2017a, b). These interactions were taken as evidence for an interplay between cognitive control and local WM content-specific processing. However, these non-invasive studies leave the mechanism by which these interactions could occur unclear – and, in particular, whether PAC at the EEG level has functional consequences at the single cell level. Here, we now provide direct evidence that PAC neurons are related to both local processing and long-range interactions, thereby providing a single-neuron mechanism for bridging these two levels of processing so commonly seen at the macroscopic level. We show that the ongoing WM content-specific processes of WM maintenance by memoranda-selective persistently active category neurons is accompanied by phase coupling to local gamma rhythms in the hippocampus. Gamma activity, in turn, was coordinated by the phase of theta activity. Crucially, single neurons that followed the local interactions between theta phase and gamma amplitude played a functional role in receiving cognitive control signals from vmPFC, reflected by stronger cross-regional theta phase coupling in trials with higher WM load and faster RT. This effect was specific to PAC neurons. Together with our noise correlation results (see below), we thus suggest that PAC neurons facilitate the temporal coordination of hippocampal processes of WM maintenance with frontal cognitive control processes.

Ultimately, cognitive-control related increased fronto-temporal coordination has to lead to increased fidelity of the retained memories. Here, we showed that this was indeed the case, albeit in a way that is only visible when examining the structure of population level activity. We show that one result of this type of cognitive control is noise correlations among cells that enhance information content. This led to pairs of simultaneously recorded PAC and category neurons with positively correlated spike counts across times within single trials. These noise correlations shaped the geometry of stimulus category information such that decodability of WM content improved. These decodability enhancements were abolished when noise correlations among neurons were destroyed. This shows that PAC neurons did not contain information about the stimulus category in isolation, which would have improved decodability even when noise correlations were removed. These findings demonstrate a functional role of PAC neurons in shaping the geometry of category representations among tuned neurons, which results in enhanced fidelity of WM memoranda representations.

Our report is, to our knowledge, the first to describe a functional role for noise correlations between neurons in humans. In particular, here we for the first time show that noise correlations among PAC and category neuron pairs predict WM-related behavior, thereby showing their behavioral relevance. This is in contrast to earlier work in macaques, which showed increased decodability of WM content but did not provide a link with WM-dependent behavior (Leavitt et al. 2017b). Of note, noise correlations are classically thought to be detrimental because they can be information limiting (Moreno-Bote et al. 2014; Bartolo et al. 2020). However, recent work reveals scenarios in which noise correlations can be beneficial (Zylberberg et al. 2016; Shahidi et al. 2019; Stefanini et al. 2020; Valente et al. 2021; Panzeri et al. 2022). We conclude that cognitive control exerted through PAC neurons can stabilize WM representations and thereby enhance the readout of WM content, leading to faster RTs. This finding suggests that noise correlations among PAC and memoranda-selective persistently active neurons might be a mechanism for stabilizing WM representations and their underlying persistent neural activity against noise or distractors. In line with this interpretation, noise correlations in our study were especially beneficial to behavior in load 3 where competing WM representations co-exist in the neural population. A hypothesis from our work that remains for further exploration is that noise correlations become stronger in the presence of distractors to enhance control over neural activity.

Taken together, our results are in agreement with a multicomponent view of WM (Cowan 2010; Baddeley 2012), where frontal control processes regulate and manage maintenance of WM content in more posterior storage-related areas (Curtis and D’Esposito 2003; Lara and Wallis 2014, 2015; D’Esposito and Postle 2015). Here, we now provide insights into the previously unknown cellular mechanisms involved in the interplay of the different components that support WM. By analyzing how single neuron activity relates to interactions of theta and gamma signals in the human brain, we provide mechanistic insights into how brain processes of cognitive control and WM content processing interact when stimulus information needs to be actively maintained through persistent activity for a short period of time. We show that ‘PAC neurons’ exist, thereby revealing a single cell correlate of the widespread macro-scale phenomena of theta-gamma PAC. The theta-gamma interactions mediated by such PAC cells play a role in cognitive control and shape WM fidelity through noise correlations with memoranda-selective persistently active neurons. PAC-mediated interareal interactions might serve as a general mechanism for top-down control to influence bottom-up processes, a hypothesis that we confirm here for WM, but which remains to be tested for other high-level cognitive functions such as attention, decision making, and LTM retrieval.

## Methods

### Patients

36 patients (44 sessions; 21 females; age: 40.47 ± 13.76 years; see Table S4) participated in the study. All patients had Behnke-Fried hybrid electrodes (AdTech Inc.) implanted for intracranial seizure monitoring and evaluation for surgical treatment of drugresistant epilepsy. Their participation was voluntary, and all patients gave their informed consent. This study was part of an NIH Brain consortium between three institutions (Cedars-Sinai Medical Center, Toronto Western Hospital, and Johns Hopkins Hospital) and approved by the Institutional Review Board of the institution at which the patient was enrolled. A pre-operative MRI together with either MRI or CT post-operative images were used to localize the electrodes as previously described (Minxha et al. 2020). Electrode positions are plotted on the CITI168 Atlas Brain (Tyszka and Pauli 2016) in MNI152 co-ordinates for the sole purpose of visualization (Fig. 1b). The 3D plot was generated using FieldTrip and the Brainnetome atlas (Oostenveld et al. 2011). Coordinates appearing in white matter or outside of the target area is due to usage of a template brain. 9 electrodes that were localized outside of the target area in native space were excluded from analysis (8 out of a total of 265 recording sites).

### Task

The task is a modified Sternberg task with a total of 140 trials and 280 novel pictures. Each trial started with a fixation cross presented for 0.9 to 1.2 s (see Fig. 1a). Depending on the load condition, the fixation cross was followed by either one (load 1; 70 trials) or three (load 3; 70 trials) consecutively presented pictures, each remaining on the screen for 2 s. In load 3 trials, pictures were separated by a blank screen randomly shown for 17 to 200 ms. Picture presentation was followed by a 2.55 to 2.85 s long maintenance period in which only the word “HOLD” was presented on the screen. The maintenance period was terminated by the presentation of a probe picture, which was either one of the pictures shown earlier in the trial (match) or a picture already presented in one of the previous trials (non-match; see below). The task was to indicate whether the probe picture matched on of the pictures shown earlier *in the same trial* or not. The probe picture was shown until patients provided their response via button press. The response mapping switched after half the trials, which was communicated to patients during a short half-time break. Responses were provided using a Cedrus response pad (RB-844; Cedrus Inc.). All pictures were novel (i.e., the patient had never seen this particular image) and were drawn from five different visual categories: faces, animals, cars (or tools depending on the version), fruits, and landscapes. Images (width x height: 10.5 x 7 visual degrees) were presented in the center of the screen and never more than twice (i.e., when serving as the probe picture). Pictures were only repeated when presented as the probe stimulus. To make sure that also the non-match probe pictures were never completely new to patients (as were the matching probe pictures), which could have been used as a strategy to solve the task without utilizing WM, we always used a picture that patients had seen already in one of the earlier trials, randomly drawn from one of the categories not used during encoding. If a patient participated in more than one session, we used a completely new set of pictures in each session to ensure that all pictures were novel in all sessions.

### Spike sorting

Each hybrid depth electrode contained eight microwires from which we recorded the broadband LFP signal between 0.1 and 8,000 Hz at a sampling rate of 32 kHz (ATLAS system, Neuralynx Inc.; Cedars-Sinai Medical Center and Toronto Western Hospital) or 30 kHz (Blackrock Neurotech Inc.; Johns Hopkins Hospital) depending on the institution. All recordings were locally referenced within each recording site by using either one of the eight available micro channels or a dedicated reference channel with lower impedance provided in the bundle, especially when all channels contained recordings of neuronal spiking. To detect and sort spikes from putative single neurons in each wire we used the semiautomated template-matching algorithm Osort (Rutishauser et al. 2006). Spikes were detected after bandpass filtering the raw signal in the 300-3,000 Hz band (see Fig. S1 for single cell quality metrics). All analysis in this paper (including the LFP) is based on signals recorded from micro wires.

### LFP preprocessing

Before analyzing the LFPs, we removed spike waveforms (action potentials) and excluded trials with inter-ictal discharges and high-amplitude noise. First, to avoid leakage of spiking activity into lower frequency ranges (Zanos et al. 2011; Anastassiou et al. 2015), we removed the waveforms of detected spikes from the raw signal by linear interpolating the raw data from -1 to 2 ms around each spike onset in the raw recording before downsampling. Since the same spike can, in rare instances, be recorded on more than one wire, we not only interpolated the data for the wire on which the neuron was detected but also for all other wires in the same wire bundle. We then lowpass filtered the raw signal using a zero phase-lag filter at 175 Hz and downsampled to 400 Hz. Line noise was then removed between 59.5 and 60.5 Hz as well as between 119.5 and 120.5 Hz using zero phase-lag band-stop filters.

Artifact and inter-ictal discharge detection was performed on a per trial and wire basis using a semiautomated algorithm together with subsequent visual inspection of the data. To detect high-amplitude noise events as well as inter-ictal discharges, we z-sored the amplitude in each channel across all trials. To avoid artifactual amplitude biasing, we first capped the data at 6 SD from the mean and re-performed the z-scoring on the capped data (see, Stark et al. 2014; Norman et al. 2019). If a single time sample in each trial and wire exceeded a threshold of 4 SD, the trial was removed from the analysis for that wire. Jumps in the signal were detected by z-scoring the difference between every fourth sample of the capped signal. Trials in which any jump exceeded a z-score of 10 SD were removed. The result of this cleaning process was visually inspected in every recording and any remaining artifactual trials were removed manually. If a wire or brain region showed excessive noise or epileptic activity, it was entirely removed from the analysis. On average, 20.4 ± 13.9 trials (14.6% of the data) were removed per wire.

### Phase-amplitude coupling

We measured the strength of phase-amplitude coupling for a wide set of frequency combinations in all recorded micro channels (except those excluded, see above) using the modulation index (MI) as introduced by (Tort et al. 2010). Since the cleaning process described above produced a different set of available trials for each channel, we first randomly sub-sampled from all correct trials in each channel such that the number of trials were the same for both load 1 and load 3. We then extracted the LFP starting at - 500 till 3,000 ms following the maintenance period onset in each selected trial. We then filtered (using pop_eegfiltnew.m from EEGLAB) (Delorme and Makeig 2004) each trial separately within the respective frequency bands of interest (see below for more details). We then extracted the instantaneous phase from the lower frequency signal and the amplitude from the higher frequency signal using the Hilbert transform. Lastly, we cut each trial to the final time window of interest of 0 – 2,500 ms relative to maintenance period onset. This last step ensures that filter artifacts that arise at the edges of the signal are removed. Next, we concatenated the phase and the amplitude signal across trials and computed the MI as described in (Tort et al. 2010) (18 phase bins). We computed MIs separately for load 1 and load 3 trials, as well as across all selected trials (both loads) to select for significant PAC channels in an unbiased fashion (see below).

To standardize the MI in each channel and condition, we computed 200 surrogate MIs by randomly combining the phase and amplitude signals from different trials (trial-shuffling), again separately for load 1, load 3, and for all trials. We fit a normal distribution to these surrogate data (normfit.m) to obtain the mean and standard deviation of each distribution. These values were then used to z-transform the raw MI values in each condition and channel. Standardizing MI values separately for each load eliminates potential systematic differences between load conditions that might arise due to load-related power or phase-differences, which could drive observed differences in PAC between conditions (Aru et al. 2015). Significant PAC channels were selected if the normalized MI computed across all selected trials (both loads) exceeded a z-score of 1.64 (p < 0.05, right-sided).

We repeated the above procedure for all frequency combinations. The phase signals were extracted for center frequencies between 2 and 14 Hz in steps of 2 Hz (2 Hz fixed bandwidth). The amplitude signals were extracted for frequencies between 30 and 150 Hz in steps of 5 Hz. The bandwidth of the amplitude signals was variable and depended on the center frequency of the low-frequency signal. It was chosen such that it constituted twice the center frequency of the phase signal (for instance, if combined with an 8 Hz center frequency for the phase signal, the bandwidth of the amplitude signal was chosen to be 16 Hz). This procedure ensures that the side peaks that arise if the amplitude signal is modulated by a lower-frequency phase signal are included (Aru et al. 2015).

### Duration of high-amplitude gamma events

To extract high-amplitude gamma events from significant PAC channels, we followed a similar rationale as described in (Norman et al. 2019). We used the same set of sub-sampled trials as used for the PAC analysis in each channel (see above). First, in each channel we extracted the raw data from each selected trial between -500 and 3,000 ms around maintenance period onset, filtered the data between 70 and 140 Hz (pop_eegfiltnew.m), and extract the instantaneous amplitude using the Hilbert transform for each trial. The data were then re-cut to 0 – 2,500 ms after the maintenance period onset, concatenated across trials within each condition and z-scored. To avoid biases introduced by extreme amplitude values, we capped the data at z-values of 3 SD and re-computed the z-transformation on the capped data (Stark et al. 2014; Norman et al. 2019). We then extracted the number and duration of events that reached a z-score of 3. The timestamps when the signal crossed a z-score of 2 were selected as the start or the end of an event, respectively. Only events that lasted at least 6 samples (15 ms, i.e., at least one full cycle of the lowest frequency of 70 Hz) were included for the analysis. Gamma event duration was then averaged across all channel and condition.

### Category cell selection

We selected for neurons whose response following stimulus onset differed systematically between the picture categories of the stimuli shown. To do so, for each trial we counted the number of spikes a neuron fired in a window between 200 to 1,000 ms after stimulus onset (all encoding periods and the probe period). We then grouped spike counts based on the category of the picture shown in that trial. For each neuron, we computed a 1×5 ANOVA with visual category as the grouping variable, followed by a posthoc one-sided t-test between the category with maximum spike count and all other categories. We classified a given neuron as a category neuron if both tests were significant (p < 0.05, 2,000 permutations (see below)). We refer to the category with the maximum firing rate as the preferred category of a cell.

### Spike-field coherence

To measure how strongly the spiking activity of a neuron followed the phase of an LFP in a certain frequency, we computed the spike-field coherence. Here, we measured SFC as the mean vector length (MVL) across spike-phases for all neuron-to-channel combinations available within a bundle or across regions (within the same hemisphere) in correct trials (Minxha and Daume 2022). To estimate the instantaneous phase from LFPs in different frequency ranges during the WM maintenance period, we first extracted data between -500 and 3,000 ms around the maintenance period onset from all clean trials in each channel and computed a Morlet wavelet convolution for 40 logarithmically spaced frequencies between 2 and 150 Hz in each trial. The trials were then cut to the final time window of interest of 0 to 2,500 ms after the maintenance period onset to remove filter artifacts at the edges of each trial. To further avoid a bias of the MVL based on differences in spike count, we subsampled spikes such that an equal number of spikes was available in each condition. We included neurons that had at least 50 spikes available in each condition (we used a minimum of 10 spikes for the preferred vs non-preferred analysis in category neurons due to a potentially low spike count in the non-preferred condition (Kamiński et al. 2020)). Next, we extracted the phase in the LFP closest to the timestamp of each spike, averaged across all spike-phases in polar space, and computed the MVL for each of the 40 frequencies. We repeated this subsampling 500 times and averaged the resulting MVLs across all repetitions within conditions. To avoid potential bias of load within the preferred vs non-preferred (category neurons, Fig. 3) or fast vs slow RT SFC comparison (cross-regional analysis, Fig. 5), we computed the SFC estimates within each load condition and then averaged across the loads.

The resulting MVL in each neuron-to-channel combination was further normalized using a surrogate distribution, which was computed after adding random noise to the timestamps of all spikes within a condition for 500 times. Potential biases of the MVL based on systematic differences between the conditions (such as power differences between conditions within a given frequency band) were thereby reduced. Like for the measure of PAC (see above), we fit a normal distribution to the surrogate data and used the mean and the standard deviation of that distribution to z-score the raw MVL within each condition.

### Selection of PAC cells

We selected for neurons whose firing rate was correlated with both theta phase and gamma amplitude during the maintenance period of the task. For all neuron-to-channel combinations within a bundle of micro wires, we extracted the data from correct trials between -500 to 3,000 ms relative to the maintenance period onset and estimated the phase of theta signals by filtering between 3 and 7 Hz and computing a Hilbert transform in each trial. Gamma amplitude was determined by computing wavelet convolutions for frequencies between 70 and 140 Hz in frequency steps of 5 Hz. Trials were cut to 0 to 2,500 ms after maintenance period onset to remove edge artifacts and then concatenated. The extracted amplitudes in each gamma frequency were z-scored across all trials and averaged across all frequencies. Computing wavelet convolutions in 5 Hz steps and z-scoring the data before averaging avoided biasing power estimates to lower frequencies due to the power law. Next, for each neuron-channel pair, we performed a median split of gamma amplitudes and binned all amplitudes into low and high gamma, respectively. In each of the two gamma groups, we further binned the corresponding theta phases into 10 groups (36° bins), resulting in a total of 20 bins (see Fig. 4a). In each of those bins, we then counted the number of spikes that occurred in each theta-gamma bin.

We fit three Poisson generalized linear models (GLMs) for each neuron-to-channel combination. In model 1, spike count (SC) was a function of theta phase (10 levels; separately as cosine and sine due to the circularity of phase values (Al-Daffaie and Khan 2017)), gamma amplitude (2 levels), and the interaction between theta phase and gamma amplitude. Model 2 included the theta phase and gamma amplitude as main effects but not the interaction term. Model 3 included a main effect for theta phase and an interaction term but no main effect for gamma amplitude:

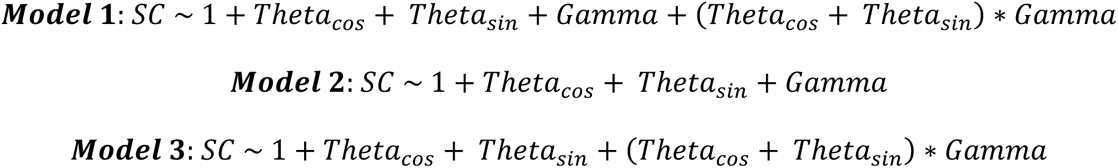

We next compared pairs of models using a likelihood-ratio test between model 1 and the two other models (using compare.m). A neuron qualified as a “PAC neuron” if model 1 explained variance in spike counts significantly better than both of the other two models (p < 0.01, FDR corrected for all possible channel combinations). The rationale behind each model comparison was the following. First, we were specifically interested in neurons that followed the interaction of theta phase and gamma amplitude, i.e., PAC, and not just theta phase or gamma amplitude alone. Selecting neurons for which model 1, including the interaction term, explained spike count variance of a given neuron significantly better than model 2, lacking the interaction term, ensured extracting those neurons. Second, we additionally compared model 1 to model 3, lacking the gamma term, for the following reason. Assume that a given neuron-channel combination has an LFP with strong PAC at the field potential level, i.e., strong interactions between theta phase and gamma amplitude, and a neuron whose firing rate is not related to neither theta phase nor gamma amplitude. Nevertheless, this situation would result in a significant interaction term in model 1 because the spikes that fall into the low and high gamma amplitude groups will have different theta phases (due to PAC). This is only the case if the underlying PAC in the LFP is very strong (see Fig. S4 for illustration and further discussion). In this scenario, however, the gamma amplitude term (nor the theta phase term) would not be significant. Comparing model 1 to model 2 *and* model 3 therefore ensures that only cells were selected at PAC cells in which the interaction term explained variance above and beyond the main effects and interactions alone.

Since at the LFP level we did not observe strong PAC in frontal regions, we restricted this analysis to channels from the MTL regions and performed this analysis separately in each load condition. If spike count variance was significantly better explained by model 1 than the two other models in either of the load conditions for at least one neuron-to-channel combination, we included this neuron as a PAC neuron. If a neuron was selected in more than one neuron-to-channel combination, we selected the combination with the highest R-squared in the full model (model 1). This combined channel was later used for within-region SFC as well as FR correlation analyses. Lastly, to determine whether the number of selected PAC neurons per area was significantly higher than chance, we repeated the entire selection process for 200 times after randomly scrambling spike timestamps and thus destroying their relationship with theta phase and gamma amplitude. The p-values indicate the proportion of repetitions that resulted in a higher number of selected neurons using the shuffled data than the original number of PAC neurons determined using the unshuffled data.

### Noise correlations and optimized population decoding

We investigated the effect of noise correlations among groups of simultaneously recorded neurons on population decoding accuracies for the image category currently hold in mind and on WM behavior during the maintenance period. To estimate noise correlations among pairs of category and PAC neurons, for each neuron we counted spikes in bins of 200 ms that slided across the maintenance period (0-2.5 s after the last picture offset) in steps of 25 ms. We then computed the correlation coefficient across all 101 time-bins in each single trial for each pair of neurons and averaged across all considered trials within each condition. We only used correct trials for this analysis, and only paired neurons that were recorded in the same session and within the same brain region. Pairs of neurons recorded on the same channel were not considered as a pre-caution against spurious correlations caused by spike sorting inaccuracies.

To investigate the contribution of PAC neurons on the population category decoding accuracy when noise correlations among neurons were intact or removed, we utilized the approach introduced by (Leavitt et al. 2017b). To measure how much a single neuron affects the decoding accuracy of an ensemble of neurons, this approach finds optimized neuron ensembles that have maximal decoding accuracy by adding each single neuron to the ensemble in a stepwise manner. Each neuron’s contribution to the ensemble can thereby be determined. In more detail, using a linear decoder, first the decoding performance for each single neuron in each region is determined from all correct trials. The neuron with the best decoding performance is then paired with each remaining neuron to determine which pair yields the best decoding accuracy. This most informative pair of neurons is then again combined with each remaining neuron to determine the most informative trio of neurons, and so on. These steps were repeated until all neurons were part of the decoding ensemble.

Since we were most interested in decoding picture category from firing rates in the maintenance period, we used trials from load 1 only. This is because the maintenance period in load 3 trials contains intermixed information about the three different categories maintained in WM. We trained a linear support vector machine (SVM) decoder (fitcecoc.m; ‘one-vs-one’) on 80% of trails and tested it on the remaining 20% using z-scored firing rates. To ensure equal amount of data for all five categories, we subsampled trials to match the lowest number of trials available in each stimulus category. Noise correlations among neurons were left intact by using the same trials for each neuron or removed by shuffling trials per neuron within each category. Shuffling trials within each category ensures that the label in each trial was still correct but the correlations among neurons were removed. We repeated each decoding analysis 500 times and averaged the results to generalize across trial selections. Since we were interested in how much a given PAC neuron contributed to the decoding performance, we tested contributions between intact and removed noise correlations only for PAC neurons that were added to the ensemble before maximal decoding performance was reached in each session and area (Leavitt et al. 2017b).

### Statistics

Throughout the manuscript, we use t-tests, ANOVAs, or mixed-model GLMs (fitglme.m in MATLAB) to assess statistical differences between conditions. T-tests and ANOVAs were calculated using permutations statistics (statcond.m as implemented in EEGLab), i.e., a non-parametric test that does not make assumptions about the underlying distributions, with 10,000 permutations unless stated otherwise. The corresponding t and F estimates, which are computed based on a normal distribution, are provided as reference only. SFC estimates tested across several frequencies were corrected for multiple comparisons using cluster-based permutation statistics as implemented in FieldTrip (Maris and Oostenveld 2007) with 10,000 permutations and an alpha level of 0.025 for each one-sided cluster, which was additionally Bonferroni corrected for the number of tests involved. Depending on whether we used z-scored firing rates or spike counts, we used mixed model GLMs based on a normal or Poisson distribution, respectively. Lastly, error bars shown in figures reflect standard errors of the mean, unless stated otherwise.

## Acknowledgments

We would like to express our deepest gratitude to the patients who volunteered to participate in this study. We thank the clinical teams at Cedars-Sinai Medical Center (in particular, Jeffrey Chung and Lisa Bateman), Toronto Western Hospital, and John’s Hopkins School of Medicine for patient management and care and support of data acquisition. We further thank Tessa Rusch and Juri Minxha for valuable discussions, Nand Chandravadia for assistance with data processing, and Ian Reucroft for assistance with data acquisition. This study was supported by a German National Academy of Sciences Leopoldina Postdoctoral fellowship (to JD), a Center for Neural Science and Medicine at Cedars-Sinai Postdoctoral fellowship (to JD), the BRAIN initiative through the National Institute of Neurological Disorders and Stroke (U01NS103792 and U01NS117839 to UR), and the National Science Foundation (BCS-2219800 to UR).

## Author Contributions

JD, JK, and UR conceived the project; JD, JK, AGPS, and YS performed experiments; JD, JK, and UK performed data analyses; CR provided patient care and supported data acquisition; TAV, WA, and ANM managed patients and performed surgeries; JD and UR wrote the manuscript with input from all authors. JD and UR acquired funding.

