## supplementary information for "Control of working memory maintenance by theta-gamma phase amplitude coupling of human hippocampal neurons"

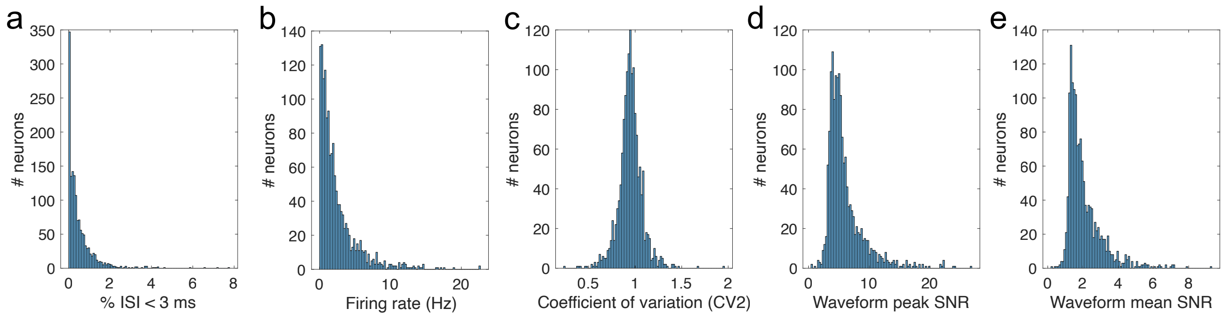


**Figure S1. Spike-quality metrics for all identified putative single units. (a)** Proportion of inter-spike intervals (ISI) below 3 ms. **(b)** Average firing rate. **(c)** Coefficient-of-variation. **(d)** Signal-to-noise ratio (SNR) for the peak of the mean waveform across all spikes as compared to the standard deviation of the background noise. **(e)** Mean SNR of the waveform.

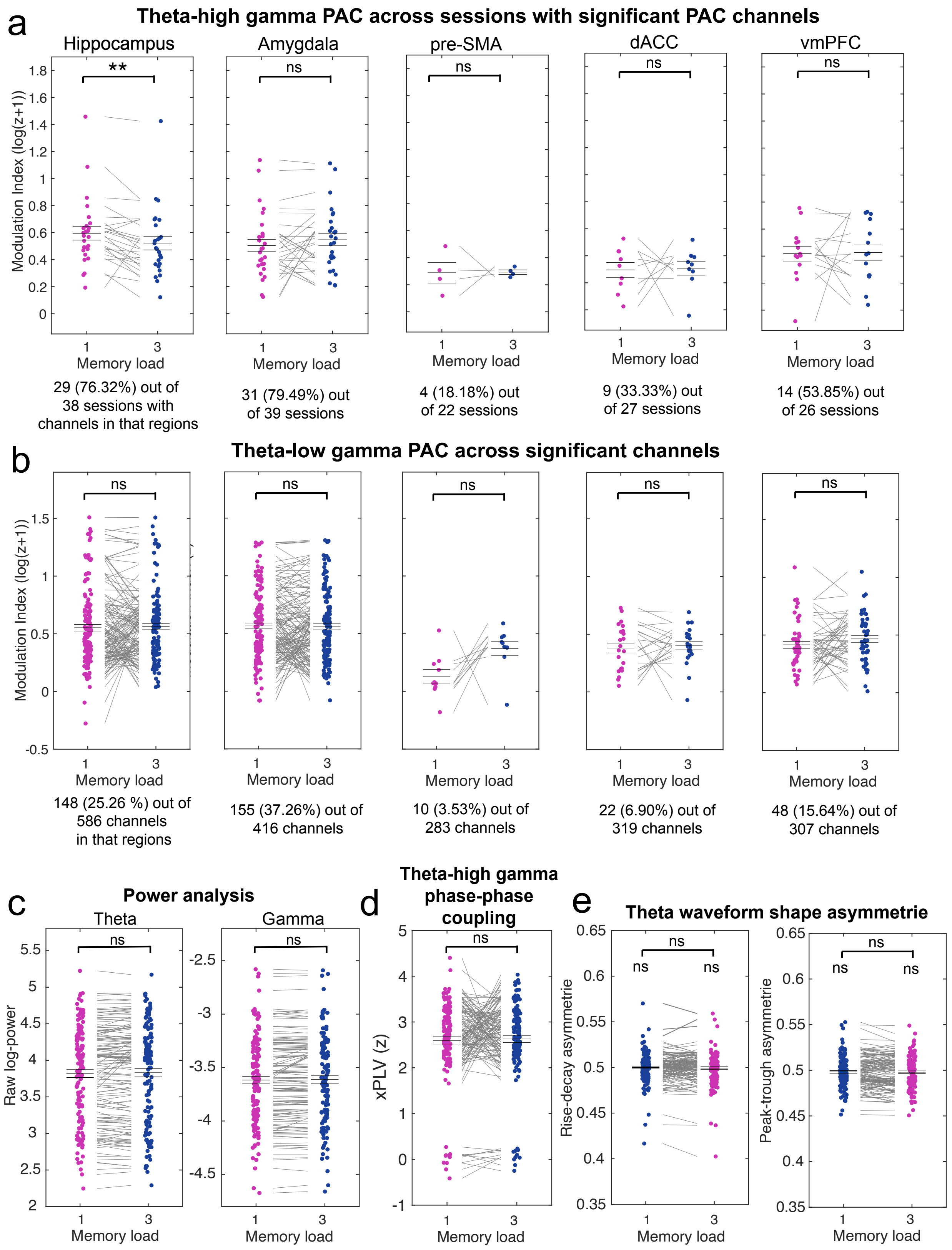
a
Theta-high gamma PAC across sessions with significant PAC channels
Hippocampus
Amygdala
pre-SMA
dACC
vmPFC
**
ns
Modulation Index (log(z+1))
Memory load
29 (76.32%) out of 38 sessions with channels in that regions
31 (79.49%) out of 39 sessions
4 (18.18%) out of 22 sessions
9 (33.33%) out of 27 sessions
14 (53.85%) out of 26 sessions
b
Theta-low gamma PAC across significant channels
148 (25.26 %) out of 586 channels in that regions
155 (37.26%) out of 416 channels
10 (3.53%) out of 283 channels
22 (6.90%) out of 319 channels
48 (15.64%) out of 307 channels
c
Power analysis
Theta
Gamma
Raw log-power
d
Theta-high gamma phase-phase coupling
xPLV (z)
e
Theta waveform shape asymmetrie
Rise-decay asymmetrie
Peak-trough asymmetrie

**Figure S2. Additional PAC analyses and controls. (a)** In addition to testing theta-gamma PAC across significant channels (see main text), we tested PAC between the load conditions across sessions after averaging all significant PAC channels within each session. The results were similar to the analysis across channels, with strongest PAC in MTL areas and only weak PAC in frontal channels (see percent of sessions with significant PAC channels below each figure), suggesting that the results were not driven by channels from a single session. Only in the hippocampus again, PAC was stronger for load 1 as compared to load 3, observable in almost each single session. No significant differences were found between the load conditions in other regions. Z-scored PAC values were shifted into a positive range by an offset of 1 and log-transformed for illustrative purposes only. All statistics are based on non-transformed z-values. **(b)** Theta to low gamma (30-55 Hz) PAC analyses. We also found strongest theta-low gamma PAC in MTL regions as opposed to frontal regions (see percentages below each figure). But for the low gamma band, we did not observe significant differences between the load conditions in any of the regions. **(c-e)** Theta-high gamma control analyses. **(c)** Comparison of theta and gamma power. The significant hippocampal PAC channels showed no differences in theta or gamma power between the two load conditions. **(d)** If the differences between the load conditions observed for PAC channels in the hippocampus were explained by waveform shape differences/theta harmonics, we should also observe an effect for cross-frequency phase-phase coupling between the same frequency bands. We tested for that in all significant hippocampal PAC channels and did not observe a significant difference. Theta-high gamma phase-phase coupling was computed as described in (Siebenhühner et al. 2020; Minxha and Daume 2022). **(e)** To further determine the influence of theta waveform shape on PAC, we tested for differences in theta waveform peak-to-trough as well as rise-to-decay asymmetries between the two load conditions which could potentially explain observed differences in theta-gamma PAC (Kramer et al. 2008; Cole and Voytek 2017). To extract and characterize each theta cycle during the delay period in all significant hippocampal PAC channels, we used the bycycle toolbox (Cole and Voytek 2019) in Python. We averaged estimates for peak-to-trough as well as rise-to-decay asymmetries across cycles during the maintenance period from the same trials used for our PAC analysis within each load and tested the estimates between the conditions. We did not find systematic differences between the conditions for both measures. Moreover, average theta waveforms were overall symmetric as both measures were not significantly different from .5 in any of the conditions. ** $p < 0.01$. permutation-based t-test.

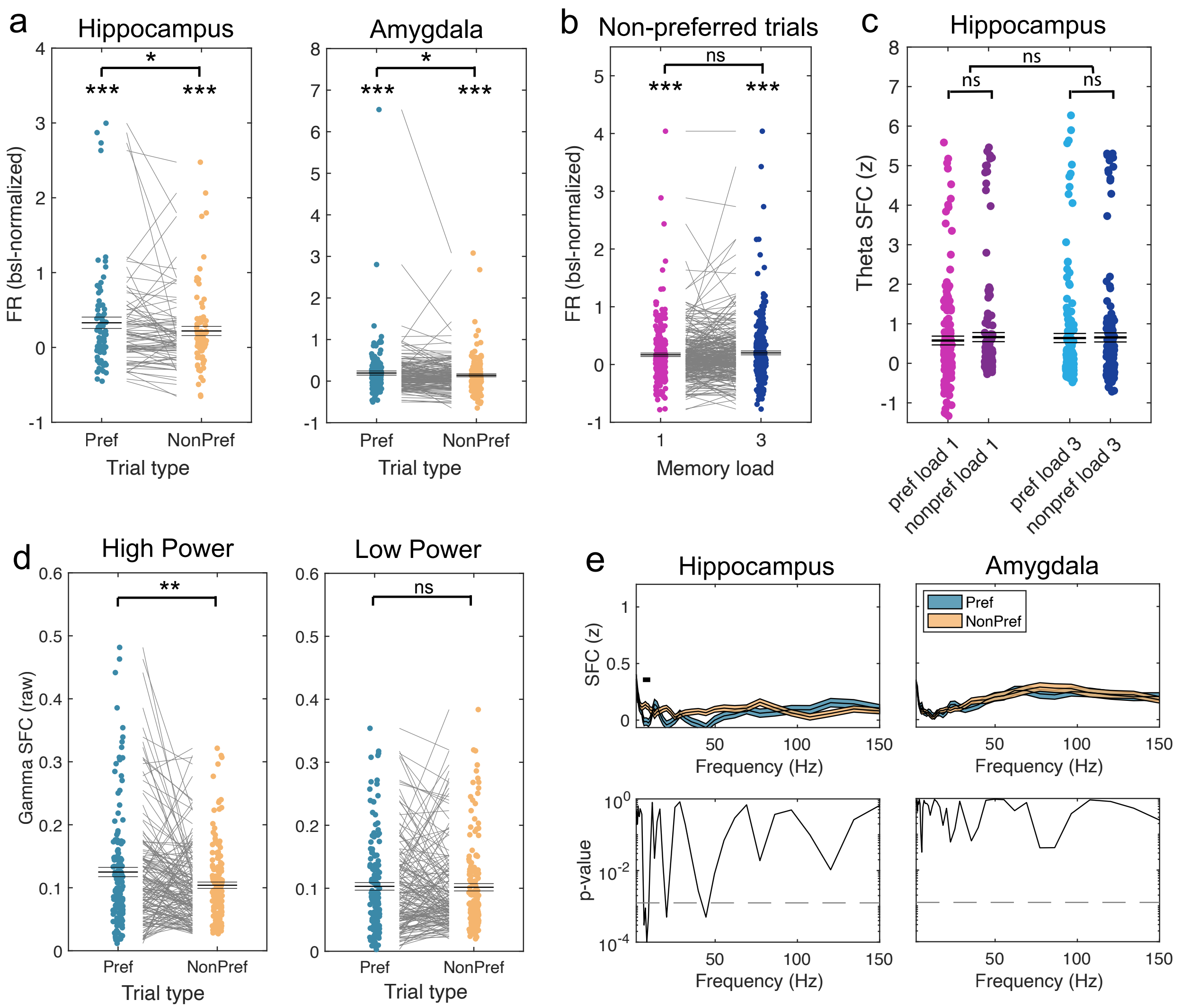


**Figure S3. Additional analyses for category cells within the MTL. (a)** Firing rates of category cells during the delay period were significantly higher for trials in which the preferred category was maintained in WM than for non-preferred categories when tested separately in both areas of the MTL. **(b)** Unlike for preferred trials, we did not observe a load effect for category cells from the MTL when non-preferred categories were maintained during the maintenance period. **(c)** When averaging theta band (3-7 Hz) SFC values for hippocampal category neurons paired with significant PAC channels, we did not observe a significant main effect for *load* or *preference* nor a significant interaction. **(d)** We performed a median split of gamma amplitudes across trials and tested gamma SFC between category cells and significant PAC channels in the hippocampus separately for spikes that occurred during high and low gamma amplitudes (spike counts were adjusted across conditions). We observed a significant difference in gamma SFC between pre-ferred and non-preferred trials only for spikes that occurred during high, not during low gamma amplitudes. **(e)** When paired with non-PAC channels, we did not observe differences in the gamma band between preferred and non-preferred trials for normalized SFC values for category cells in hippocampus or amygdala. In the hippocampus, we observed a significant difference in the alpha range (7-11 Hz) with SFC for non-preferred trials higher than for preferred trials, which we did not further consider in our analyses. **(d)**. *** $p < 0.001$; * $p < 0.05$; ns = not significant; permutation-based t-test.

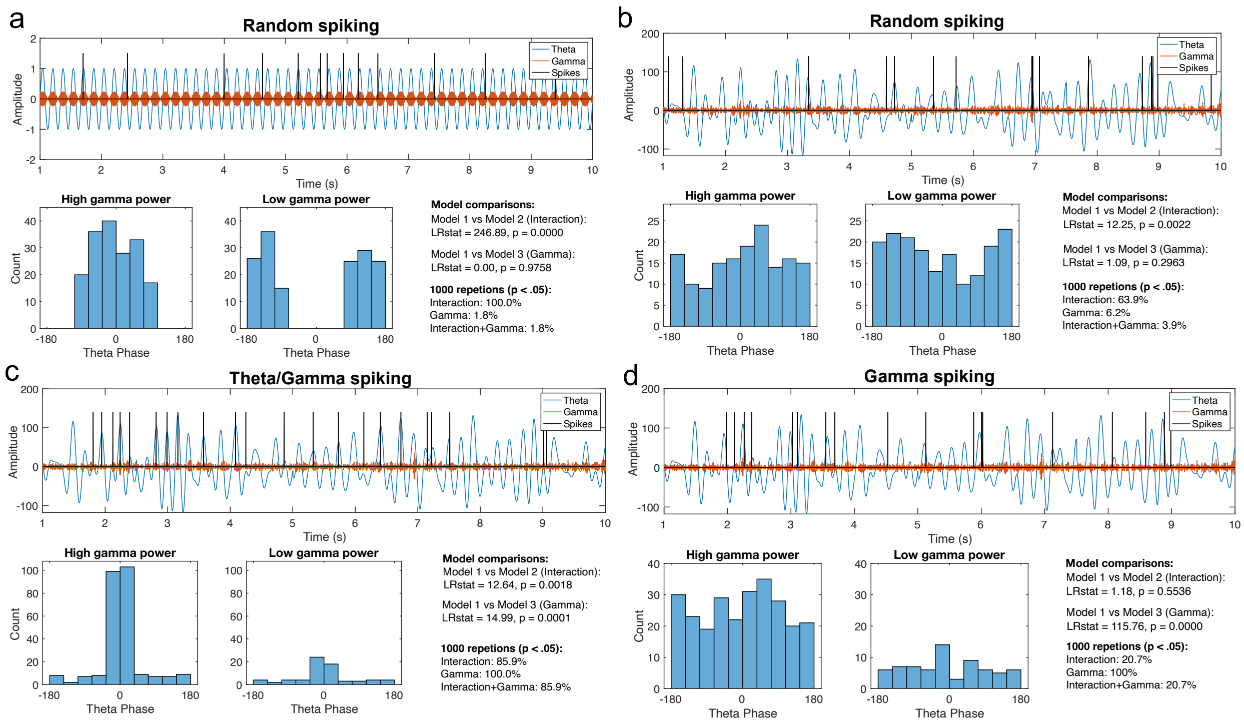


**Figure S4. Simulations supporting the PAC neuron selection approach.** We argue that a neuron that fires randomly with respect to theta phase and gamma power could still be selected as a PAC neuron if only the GLM interaction term between theta phase and gamma amplitude is considered. In addition to selecting neurons whose FRs are better explained by a model including an interaction term as compared to a model with no interaction term, we therefore introduced a second criterion by comparing the full model against a model that lacks the gamma amplitude term. The simulations presented here are meant to visualize our reasoning. In (a), we simulate theta (6 Hz) and gamma (80 Hz) signals, where gamma amplitudes perfectly couple to theta phase. This highly artificial LFP signal only serves to simplify visualization. We also include illustrations in (b) to (d) using an originally recorded LFP channel from our dataset (filtered between 3-7 Hz and 70-140 Hz) that shows strong levels of PAC. This is to show that the same arguments also hold for real data. For the purpose of these illustrations, we used an LFP signal of roughly 160 s length and simulated 300 spike timestamps (black ticks), of which 9 s are plotted. **(a)** In this simulation, we modelled random spike timestamps with respect to theta phase and gamma amplitude (upper panel). According to our GLM selection approach, we grouped spikes in 10 theta phase bins and 2 gamma amplitude bins and determined spike counts in each bin (lower panels). As can be seen from the histograms in the lower panels, the theta phase distribution of spike counts differs between low and high gamma amplitudes, resulting in a highly significant interaction term between theta phase and gamma amplitude. The reason for this is that gamma amplitude itself is already perfectly coupled to theta phase. Separating spikes into low and high gamma will therefore also result in different theta distributions among the two spike count groups. Thus, when testing a model that contains theta phase and gamma amplitudes as well as their interaction against a model without the interaction, spike counts will be highly significantly better explained by the full model, as was the case in this example ($p < 0.001$; see likelihood-ratio test results on the right). However, since the time stamps are random, we should not observe a difference in overall spike counts between low and high gamma amplitudes, which was also the case in this example ($p = 0.98$). Introducing such a gamma term comparison as a second selection criterion thus ensures that this simulated random neuron would not have been selected. In 1000 repetitions of this simulation, our approach would have selected only 1.8% of such randomly spiking neurons (see text on right side). **(b)** Similar to (a) but using a real LFP recording from our dataset that shows strong levels of PAC. 300 spike timestamps were again modelled randomly with respect to theta phase and gamma amplitude. Similar albeit weaker statistics were observed in these simulations. **(c)** Using the same LFP as in (b) but now simulating 300 spike timestamps that prefer high gamma amplitudes and a theta phase of 0 (i.e., PAC spiking plus 10% noise). Here, as desired, the full model explains spike counts significantly better than both the other models and this neuron would be selected as a PAC neuron. **(d)** In this example, we simulate a "gamma neuron", i.e., a neuron whose FR follows gamma amplitude, but not theta phase. In most cases (79.3% of 1000 repetitions), these gamma neurons were successfully rejected. Since we did not control for theta phase in these simulations using a strong LFP channel, however, around 20% of the simulations modelled PAC rather than pure gamma spiking.

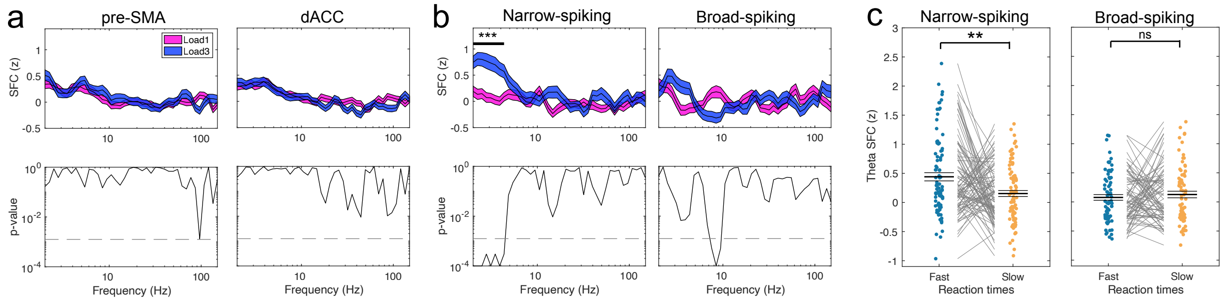


**Figure S5. Cross-regional SFC effects for pre-SMA and dACC, as well as for fast- and broad-spiking PAC neurons from the hippocampus. (a)** Cross-regional SFC between hippocampal PAC neurons and LFPs recorded in pre-SMA (left) or dACC (right) did not reveal any difference between the two WM load conditions in any of the frequencies. **(b)** Earlier work has suggested that cognitive control might especially be governed through long-range connections between frontal and sensory regions that target inhibitory interneurons to (dis-)inhibit local circuitries (Zhang et al. 2014; Hattori et al. 2017; Malik et al. 2022). We thus asked if we observe a differential effect for the hippocampal PAC neuron connections after separating them into narrow- and broad-spiking neurons based off their waveform shapes, which has been suggested to categorize neurons into inhibitory and excitatory neurons, respectively (Barthó et al. 2004; Mosher et al. 2020). For analysis of connections involving narrow- and broad-spiking PAC neurons separately, we observed a significant difference in theta SFC between load 3 and load 1 only for the narrow-spiking PAC neurons (trough-to-peak time $< 0.5$ ms; 91 connections; cluster-$p < 0.001$, left). No effect was found for broad-spiking PAC neurons (84 connections; all cluster-$p > 0.025$) **(c)** Similarly, theta SFC for fast RT was significantly stronger than for slow RT only for narrow-spiking ($t(90) = 3.02$, $p = 0.003$, left), not for broad-spiking PAC neuron connections between hippocampus and vmPFC ($t(75) = -0.66$, $p = 0.52$; right; spikes were median split into fast and slow RT trials per load condition and then averaged across loads to avoid potential confounds). *** $p < 0.001$; ** $p < 0.01$; ns = not significant; (cluster-)permutation-based t-test.

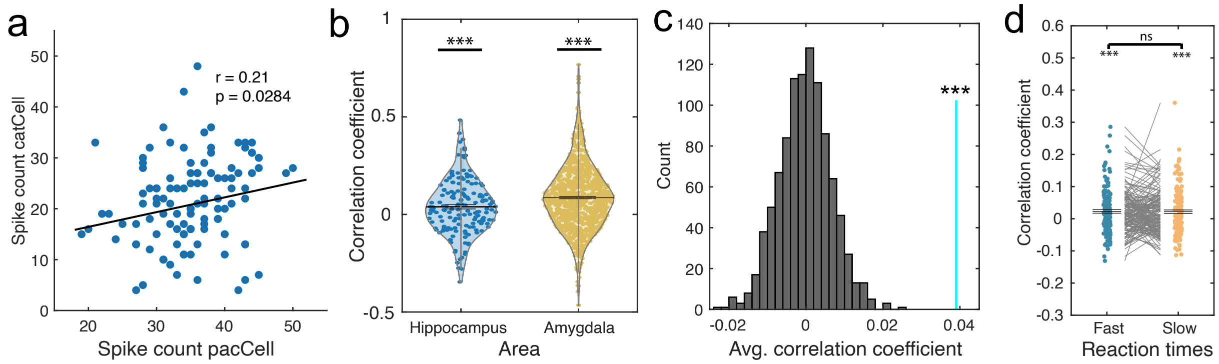


**Figure S6. Trial-by-trial noise correlations among PAC and category neurons.** Related to Figure 6. **(a)** Example showing noise correlations computed across trials for a pair of simultaneously recorded category and PAC neurons from the hippocampus. Each dot represents the spike count in a correct trial during the maintenance period for each neuron. For this example, the firing rate of the two neurons was positively correlated across trials. **(b)** Correlation coefficients for all possible pairs of category and PAC neurons in the hippocampus and amygdala. In both regions, correlation coefficients covered a broad range of both positive and negative values and were significantly higher than zero on average. **(c)** Repeat of the correlation analysis for all possible PAC-category neuron pairs in the hippocampus. Shuffling trial labels for 1000 times resulted in far lower correlations between pairs of neurons than unshuffled trial labels (cyan line; mean of correlation coefficients across all pairs), showing that trial shuffling successfully removed noise correlations. **(d)** Bin-wise correlations (averaged across trials) among pairs of hippocampal PAC and category neurons did not differ between fast and slow RT trials for non-preferred trials. Each dot represents the correlations coefficient for a pair after averaging, computed per trial and then averaged across all considered trials. *** $p < 0.001$; ns = not significant; permutation-based t-test.

RT ~ 1 + PAC + load + PAC * load + (1 | channelID)

| | Hippocampus (137 levels) | | | Amygdala (130 levels) | | |
|---|---|---|---|---|---|---|
| **Name** | **Estimate** | **t** | **p** | **Estimate** | **t** | **p** |
| **Intercept** | 0.38 | 7.76 | <0.001 | 0.25 | 7.68 | <0.001 |
| **PAC** | -49.07 | -4.01 | <0.001 | -5.15 | -1.02 | 0.48 |
| **Load1** | -0.48 | -7.10 | <0.001 | -0.48 | -9.91 | <0.001 |
| **PAC : Load1** | 26.30 | 1.58 | 0.11 | 5.67 | 0.52 | 0.60 |

**Table S1. GLM results for trial-by-trial correlations between RT and PAC.** Related to Figure 2. Load 1 are tested against load 3 trials.

PAC ~ 1 + $SC_{catCell}$ + load + $SC_{catCell}$ * load + (1 | neuron-channelID)

| | Hippocampus (151 levels) | | | Amygdala (423 levels) | | |
|---|---|---|---|---|---|---|
| **Name** | **Estimate** | **t** | **p** | **Estimate** | **t** | **p** |
| **Intercept** | -0.044 | -3.14 | 0.002 | 0.003 | 0.33 | 0.74 |
| **SC** | 0.002 | 2.40 | 0.017 | -0.001 | -0.75 | 0.45 |
| **Load1** | 0.076 | 3.85 | <0.001 | -0.010 | -0.76 | 0.45 |
| **SC : Load1** | -0.004 | -2.57 | 0.010 | 0.002 | 1.48 | 0.14 |

**Table S2. GLM results for trial-by-trial correlations between PAC and spike counts of category neurons.** Tested across all neuron-to-channel combinations involving significant PAC channels. Related to Figure 3. Load 1 are tested against load 3 trials. SC = spike count.

$$PAC \sim 1 + SC_{pacCell} + load + SC_{pacCell} * load + (1 \mid neuron\text{-}channelID)$$

| | Hippocampus (79 levels) | | | Amygdala (163 levels) | | |
|---|---|---|---|---|---|---|
| **Name** | **Estimate** | **t** | **p** | **Estimate** | **t** | **p** |
| **Intercept** | -0.0565 | -2.76 | 0.005 | 0.021 | 1.37 | 0.17 |
| **SC** | 0.0032 | 2.19 | 0.028 | $-3 \times 10^{-5}$ | -0.02 | 0.98 |
| **Load1** | 0.0931 | 3.26 | 0.001 | -0.054 | -2.55 | 0.01 |
| **SC : Load1** | -0.0043 | -2.21 | 0.027 | 0.001 | 0.89 | 0.37 |

**Table S3. GLM results for trial-by-trial correlations between PAC and spike counts of PAC neurons.** Related to Figure 4. Load 1 are tested against load 3 trials. SC = spike count.

| Session | Gender | Age | Seizure onset zone | Hippo | Amy | preSMA | dACC | vmPFC |
|---|---|---|---|---|---|---|---|---|
| **P55cs** | f | 43 | Right mesial temporal | 1/14 | 21/14 | 12/14 | 10/14 | 2/14 |
| **P55cs_2** | - | - | - | 8/13 | 18/14 | 11/14 | 1/14 | 3/14 |
| **P56cs** | m | 48 | Bilateral mesial temporal + orbitofrontal | 4/12 | 21/13 | 5/6 | 4/14 | 8/14 |
| **P58cs** | f | 32 | Right frontal neocortical | 1/14 | 12/14 | 20/14 | 12/14 | 8/13 |
| **P60cs** | m | 67 | Left mesial temporal | 8/14 | 13/14 | 11/14 | 13/13 | 6/14 |
| **P61cs** | f | 52 | Left mesial temporal | 6/7 | 6/14 | 13/14 | 9/14 | 5/14 |
| **P61cs_2** | - | - | - | 2/7 | 8/14 | 18/12 | 0/7 | 8/13 |
| **P62cs** | f | 25 | Left mesial temporal | 0/0 | 12/7 | 7/7 | 0/6 | 5/7 |
| **P64cs** | f | 63 | Right lateral temporal neocortical | 0/7 | 17/6 | 25/14 | 47/12 | 23/13 |
| **P65cs** | f | 55 | Bilateral independent temporal | 5/0 | 18/7 | 2/14 | 1/14 | 7/7 |
| **P67cs** | f | 38 | Bilateral mesial temporal | 0/14 | 12/10 | 1/14 | 0/7 | 8/14 |
| **P68cs** | m | 54 | Bilateral mesial temporal | 16/0 | 7/3 | 7/14 | 2/13 | 4/14 |
| **P69cs** | f | 41 | Not localized | 8/15 | 7/14 | 0/7 | 10/14 | 11/15 |
| **P70cs** | f | 30 | Right temporal | 1/7 | 14/7 | 9/13 | 0/14 | 0/14 |
| **P70cs_2** | - | - | - | 1/7 | 5/6 | 5/14 | 0/14 | 1/12 |
| **P71cs** | m | 40 | Not localized | 0/14 | 3/14 | 3/14 | 10/14 | 11/14 |
| **P72cs** | f | 25 | Not localized, bilateral independent | 4/0 | 0/7 | 3/14 | 5/14 | 1/14 |

| | | | | | | | | |
|---|---|---|---|---|---|---|---|---|
| **P73cs** | f | 58 | Left mesial temporal | 5/14 | 17/15 | 4/14 | 6/14 | 9/14 |
| **P76cs** | f | 24 | Not localized | 6/14 | 24/15 | 7/14 | 16/14 | 10/7 |
| **P77cs** | f | 46 | Right auditory cortex | 4/14 | 41/15 | 9/14 | 0/0 | 26/14 |
| **P78cs** | f | 54 | Right anterior temporal | 13/7 | 14/7 | 0/0 | 0/0 | 0/0 |
| **P79cs** | f | 42 | Right anterior lateral temporal neocortex | 17/7 | 28/8 | 20/14 | 12/14 | 18/16 |
| **P79cs_2** | - | - | - | 13/7 | 19/8 | 12/14 | 16/14 | 12/15 |
| **P88T** | m | 26 | Right mesial temporal | 22/0 | 6/0 | 0/0 | 0/0 | 0/0 |
| **P89T** | f | 45 | Right frontal | 22/28 | 18/14 | 0/0 | 0/14 | 0/0 |
| **P90T** | m | 20 | Occipital cortex | 13/17 | 29/7 | 0/0 | 0/0 | 0/0 |
| **P90T_2** | - | - | - | 11/27 | 2/11 | 0/0 | 0/0 | 0/0 |
| **P90T_3** | - | - | - | 13/27 | 0/11 | 0/0 | 0/0 | 0/0 |
| **P91T** | f | 59 | Left cingulate cortex + insula + orbitofrontal | 5/13 | 1/6 | 0/0 | 0/7 | 12/7 |
| **P93T** | m | 23 | Left mesial temporal | 4/7 | 3/0 | 0/0 | 0/7 | 0/0 |
| **P96T** | f | 58 | Bilateral mesial temporal | 6/0 | 5/14 | 0/0 | 0/0 | 0/0 |
| **P101T** | f | 25 | Left neocortical temporal | 16/28 | 5/8 | 0/0 | 0/0 | 0/0 |
| **P101T_2** | - | - | - | 8/27 | 6/8 | 0/0 | 0/0 | 0/0 |
| **P103T** | m | 49 | Right mesial temporal | 8/14 | 4/6 | 0/0 | 0/2 | 4/7 |
| **P106T** | m | 26 | Multifocal | 7/21 | 14/7 | 0/0 | 0/0 | 0/0 |
| **P109T** | m | 28 | Multifocal | 15/28 | 7/14 | 0/0 | 5/13 | 4/7 |
| **P110T** | m | 38 | Right fusiform cortex | 6/19 | 10/14 | 0/0 | 0/0 | 0/0 |
| **P113T** | m | 36 | Right mesial temporal | 13/28 | 19/14 | 0/0 | 0/0 | 0/0 |
| **P113T_2** | - | - | - | 11/21 | 7/14 | 0/0 | 0/0 | 0/0 |
| **P116T** | m | 28 | Left amygdala | 9/13 | 18/13 | 0/0 | 9/14 | 0/7 |
| **P129T** | f | 25 | Bilateral mesial temporal | 25/28 | 5/14 | 0/0 | 0/0 | 0/0 |
| **P1802jh** | m | 62 | Right mesial temporal | 12/16 | 0/0 | 0/0 | 0/0 | 0/0 |
| **P1809jh** | m | 45 | Left inferior + middle frontal gyrus | 1/8 | 0/0 | 0/0 | 0/0 | 0/0 |
| **P1811jh** | f | 27 | Bilateral mesial temporal | 10/8 | 0/0 | 0/0 | 0/0 | 0/0 |

**Table S4. Patient demographics and neuron/channel count per area.** For each area, the first number represents the neuron count, the second the number of clean micro LFP channels.
